# Neurodegeneration-inducing macromolecules exit the brain via nanovascular conduits formed by reticular fibroblasts

**DOI:** 10.64898/2026.09.21.752863

**Authors:** Ross Nortley, Silvia Anderle, George Sideris-Lampretsas, Harvey Davis, Huma Sethi, Delan N. Alasaadi, Sophie Acton, David Attwell

**Author notes:** Before publication: Send correspondence to D Attwell. After publication: Send correspondence to R Nortley or D Attwell.

## Abstract

Accumulation of proteins such as amyloid β (Aβ), hyperphosphorylated tau and α-synuclein within the brain alters neural information processing and causes neurodegeneration^1–3^, but how toxic solutes are cleared from the brain remains highly controversial^4,5^. Proposed exit routes include efflux across endothelial cells into the blood^6,7^, and movement to the pial surface via vasomotion-induced pumping along spaces within arteriolar smooth muscle^8^ or via outflow along the perivascular space of ascending venules promoted by water flux through astrocytes (the glymphatic system^9^). From the pial surface of the brain, drainage may continue to dural lymphatics, along the outer sheaths of exiting cranial nerves and across the cribriform plate^10–14^. We now report the presence, in mice and humans, of 2 μm diameter conduits that remove fluorescently labelled tau and Aβ from the brain. These conduits form a spatially-organised mesh within the walls of penetrating arterioles and pial arteries, and around the surface of ascending venules and deep cerebral and pial veins. They course through the pial and arachnoid layers to span the CSF space, wrapping the brain and cranial nerves. They are formed of reticular fibroblasts, which label for VE-cadherin^15^ and PDGFRα^16^, the lymphatic markers^17^ podoplanin, VEGFR3 and Prox1, and reticular fibroblast extracellular matrix components collagen I and VI^16,18–20^. Parenchymal tau drains from the brain at a similar rate via arteriolar conduits and via conduits around venules, arguing against preferential removal by a glymphatic mechanism. In Alzheimer’s disease model mice, Aβ is seen traversing these lymph node-like conduits. Modulation of molecular transfer via this route may accelerate or delay cognitive decline, and slowed transfer from arteriolar to pial-arachnoid conduits may initiate cerebral amyloid angiopathy.

## Introduction

The accumulation of proteins, such as Aβ and hyperphosphorylated tau (p-tau), within the brain alters cognition and leads to neurodegeneration^1^. Speeding the mechanisms by which toxic solutes are cleared from the brain could be a therapeutic approach to preventing this dysfunction, however these mechanisms are poorly understood^4,5^. The outflow of cerebrospinal and interstitial fluid (CSF/ISF) is mainly via lymphatic pathways^10^ but it remains uncertain to what extent molecules known to induce dementia (such as Aβ and p-tau^1^) follow this route, rather than leaving the brain across endothelial cells of the brain vasculature^6,7^. Endogenous neuronally-derived proteins entering the brain parenchyma have been reported to follow a different exit route to proteins injected into the CSF^21^.

Clearance routes from the brain are shown in Extended Data Fig. 1a. Half of the CSF outflow drains to cervical lymphatics and deep cervical lymph nodes (dCLN), while the other half drains to mediastinal, iliac and sacral lymph nodes from the spinal cord^10,13,22,23^. Flow to cervical lymphatics/dCLN occurs via initial drainage to intracranial dural lymphatics^11–14^ perhaps via arachnoid granulations^24^, or via extracranial lymphatics by passing across the cribriform plate and along exiting cranial nerves^4^. The dura, one of three meningeal layers that envelop the central nervous system (CNS), adheres to the skull and is separated from the CSF space beneath by the arachnoid barrier cell layer (ABCL, which prevents CSF from flowing directly into the dura), as well as the inner arachnoid. The pial layer of the meninges forms the floor of the CSF space, covering the CNS parenchyma, and is connected to the inner arachnoid by trabeculae or cell processes that span the subarachnoid space^25^. The dural lymphatics are located along the dorsal brain surface, the basolateral skull regions, the anterior and middle cranial fossae, the outer dural sheaths of exiting cranial nerves (including the optic and trigeminal nerves) and around the skull base foramina.^4,11,12,14,26,27^.

Fluid and solute efflux from the brain parenchyma may occur across endothelial cells (Extended Data Fig 1b), along the smooth muscle basement membrane of penetrating arterioles and pial arteries (driven by arterial smooth muscle cell contraction - intramural periarterial drainage, IPAD, Extended Data Fig. 1c)^28,29^ or along the perivascular space of ascending venules and cerebral veins (driven by water flux through astrocytes - the glymphatic hypothesis, Extended Data Fig. 1d^9^). Onward drainage to the dural lymphatic pathways requires fluid and solutes to traverse the pia, inner arachnoid and ABCL^30^. Discontinuities within the ABCL, where bridging veins cross the CSF space from the pia to the dura, termed arachnoid cuff exit (ACE) points^31^, provide a potential route for CSF/ISF and proteins to reach the dural lymphatics. Similarly, CSF and solutes can drain through openings in the ABCL to extracranial submucosal lymphatic vessels around the cribriform plate, and the ABCL surrounding cranial nerves possesses breaches that may allow clearance to dural lymphatics surrounding the nerve or continued perineural outflow to extracranial lymphatic networks^4,32^. The cellular basis of all these exit pathways remains undefined.

Within lymph nodes and secondary lymphoid organs, immunologically-specialised myofibroblasts – termed fibroblastic reticular cells (FRCs) – form stellate cell-cell connections to create a three-dimensional lattice on which immune cells migrate^16,20^. FRCs also produce and ensheathe an organised meshwork of extracellular matrix (ECM) components, creating a conduit network (with a collagen fibre core: Extended.Data Fig. 1f) that rapidly transports lymphatic fluid and solutes, including antigens and inflammatory mediators, deep into the lymph node parenchyma^16,18–20^. Similar lymphatic drainage conduits exist within arachnoid granulations (AG)^24^ and connecting interstitial spaces within organs (skin and colon), and outside of organs (skin into subcutaneous fascia and colon into mesentery)^33^. This raises the question of whether similar conduits promote the efflux of fluid and solutes from the brain.

We now show, in wild-type (WT), NG2 Ds-Red, Pdgfra-mGFP-CreERT2 and Alzheimer’s disease model (APP^NL-G-F^) mice^34^, as well as live human tissue, that FRCs (that label for VE-cadherin, podoplanin^16,20^ and other lymphatic markers^17^) construct a spatially organised, web-like conduit system surrounding and invaginating the CNS. This lymph-node-like system transports solutes, including monomeric tau-441 and Aβ_1-42_, as well as endogenously produced mutant human^34^ Aβ, out of the brain (in the walls of penetrating arterioles and around ascending venules) and along pial veins and exiting cranial nerves.

## Results

To examine the anatomical and molecular substrates for macromolecule transport out of the brain, we performed in vivo two-photon imaging of the somatosensory cortex and/or confocal imaging of antibody-labelled fixed brain and cranial nerve tissue, following application of fluorescently tagged monomeric tau-441 and monomeric Aβ_1-42_ (chosen for their role in dementia and neurodegeneration, because efflux routes may depend on the properties of the particular molecule studied). Application was by four different routes: intracortical injection, intra-cisterna magna (ICM) injection, topical application to the CSF space through a cranial window, or live tissue incubation, and we also examined exit of endogenously produced Aβ_1-42_ in the APP^NL-G-F^ mouse. Fluorescently-labelled monomeric tau was used as the primary tracer because it gave the brightest signal, but fluorescently-tagged exogenous Aβ and endogenous Aβ followed the same route (see below). Visualising the exit pathway for tau and Aβ allowed us to define the cellular components of this pathway, as described below.

### Parenchymal tau exits the brain equally via arteriole and venule conduits

To identify efflux routes for neurotoxic proteins from the brain, we first used two-photon imaging through a cranial window (with the dura removed) following micro-injection of 500 nl monomeric fluorescent tau-441 (10 µM, at a rate of 100 nl/min^11^, half the rate used previously^14^) at a depth of 250 µm in the somatosensory cortex of WT, NG2-dsRed mice (to label mural cells) and Pdgfra-mGFP-CreERT2 mice (to label fibroblasts). Baseline imaging confirmed that no fluorescence was present in the cortex or pial-arachnoid layers before beginning the infusion (Extended Data Fig. 2a). Within minutes of starting the injection, fluorescent tau-441 was seen diffusely within the brain parenchyma, and in conduits within the smooth muscle layer of penetrating arterioles and surrounding the surface of ascending venules (Fig. 1a-b), and within the pia in conduits in arterial walls, around pial veins and crossing the surface of the brain (Fig. 1c-d). At the deep end of the arteriolar conduits, tau could be seen accumulating into conduits around capillaries (Extended Data Fig. 2b), consistent with previous reports^28^. Within the conduits, the fluorescence intensity was higher than in the surrounding extracellular space, suggesting active accumulation into conduits. In penetrating and pial arteries, the tau moved in conduits that appeared to be embedded in the smooth muscle cell walls of the vessels (Fig. 1e). By 90 min post-injection the labelled tau had disappeared from the parenchyma and conduits below the brain surface, but was still visible in some pial conduits, and remained in perivascular macrophages in the pial layer (Fig. 1f-i).

**Fig. 1.**
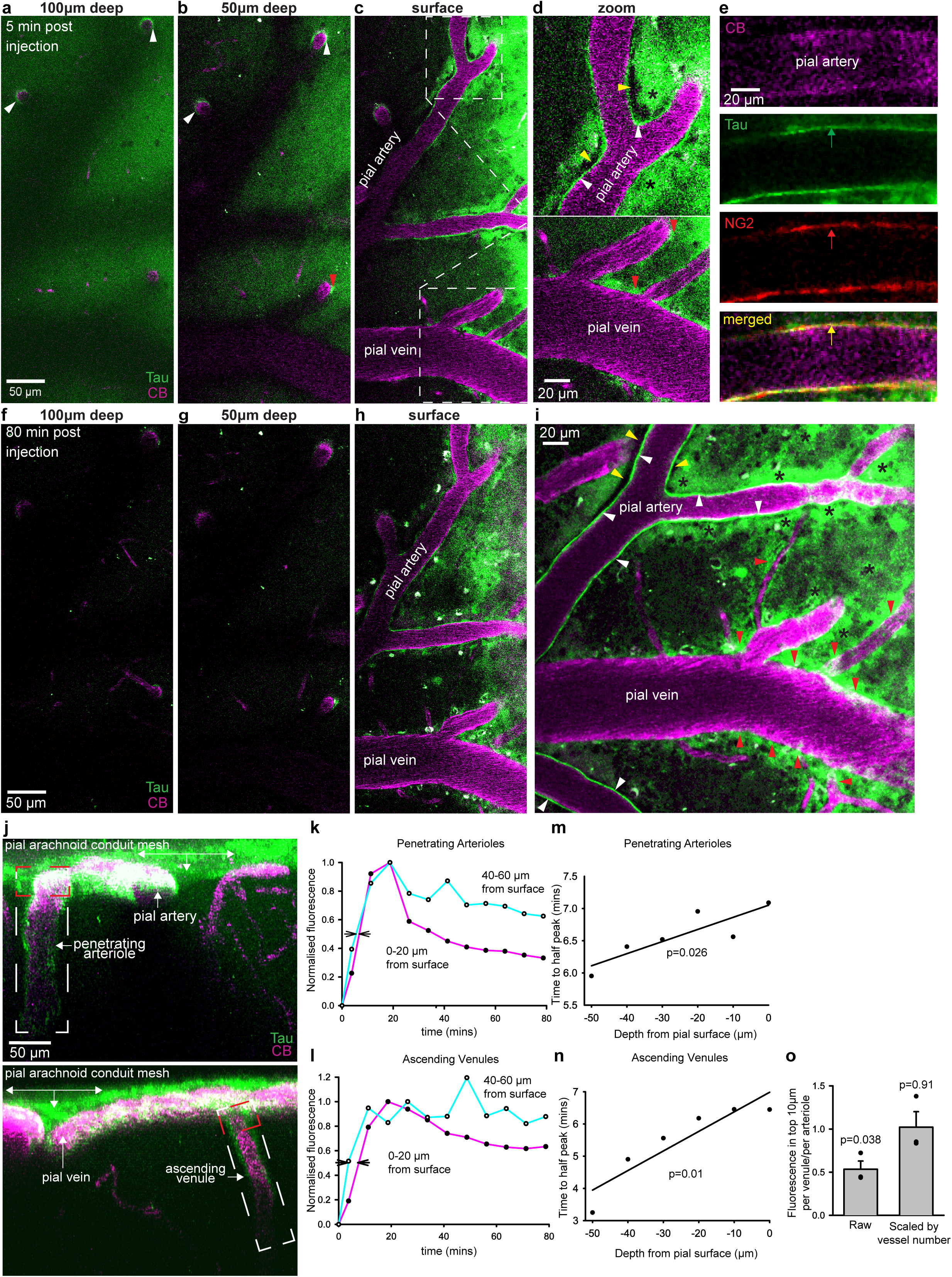
Tau exits the brain via conduits in penetrating arteriole walls and around ascending venules to enter a pial-arachnoid conduit mesh. **I**n vivo 2-photon microscopy of 8-week WT mouse cortex 5-min post parenchymal-injection of fluorescent tau-441 250 µm below the brain surface (CB, cascade blue in blood). (**a, b**) At 100 (**a)** and 50 (**b**) µm deep, tau-441 is seen diffusely in the parenchyma and more brightly in the walls of penetrating arterioles (white arrows) and around ascending venules (red arrow). (**c**) At the brain surface, tau-441 drains within pial arterial walls and around pial veins, to enter a pial-arachnoid conduit mesh). (**d**) Zoom of areas in (**c**) showing tau-441 draining in conduits in pial artery wall (white arrowheads, surrounded by an unlabelled fluid-filled space: yellow arrowheads) and around ascending venules and pial veins (red arrow heads) to enter the pial-arachnoid mesh (black asterisks). (**e**) Colocalisation of tau-441 with NG2 labelling (in NG2-DsRed mice) confirms clearance via conduits in the smooth muscle layer of pial arteries. (**f-h**) Regions in (a-c) imaged at 90 min post-parenchyma injection show tau-441 has cleared from the parenchyma, from conduits in the walls of penetrating arterioles and pial arteries, from conduits around ascending venules and pial veins, and from most of the pial-arachnoid conduit mesh, leaving residual tau in perivascular macrophages. (**i**) Larger view of the area in h. **(j)** Lateral view images of tau-441 draining from the brain within the penetrating arteriole wall (top image, white outline) and along the outer surface of an ascending venule (bottom image) to enter the pial-arachnoid conduit mesh. (**k**, **l**) Tau fluorescence 40-60 μm and 0-20 μm (red box in (j)) from the pial surface as a function of time after injection, in conduits within penetrating arterioles (**k**) and around ascending venules (**l**) in 2 animals (total 8 arterioles and 12 venules). (**m**, **n**). Half time to peak fluorescence as a function of depth in arterioles (**m**) and venules (**n**); tau arrives earlier at greater depths, implying upward transfer out of the brain. (**o**) Left: ratio of tau fluorescence in top 10 μm of ascending venules to that in penetrating arterioles, at earliest stack imaged in 3 mice. Right: raw data corrected for relative number of venules and arterioles. P values compare with 1.

Measuring the tracer intensity in three animals, in the conduits within the walls of 12 penetrating arterioles and around 20 ascending venules (Fig. 1j), showed that, in both arteriole wall and peri-venule conduits, the tracer concentration rose rapidly (within 5 mins) after injection as tracer entered the conduits, and then decayed over the subsequent 90 mins as tracer was cleared from the conduits. In 8 arteriole conduits (Fig. 1k) and 12 venule conduits (Fig. 1l) in two animals in which it was possible to image sufficiently rapidly after injecting the fluorescent tau, the time to half the maximum fluorescence was significantly faster in the lower regions of the conduits than where the arterioles and venules met the pial surface (Fig. 1m, n; p=0.026 and p=0.01 respectively), consistent with the tau moving up the conduits towards the pial surface. The decay of fluorescence was slower in both arteriole and venule conduits at greater depths in the cortex (Fig. 1k, l). The slopes of the linear regressions in Fig. 1m-n correspond to speeds of movement of tau of 0.89 μm/s in arteriole conduits and 0.27 μm/s in venule conduits, similar to the speed (0.58 μm/s) estimated for lymph node conduits^20^. The >3-fold higher speed for movement in the arteriole conduits may reflect their visibly more linear organisation within the arteriole wall (compared to the conduits around ascending venules) or greater pulsatility-driven flow from movement of the adjacent arteriole^8^ (Fig. 1d).

In 3 animals, per vessel, the amount of tracer detected in the surface 10 µm of the conduits around ascending venules in the first image stack taken after injection was only 53.5±9.4% of the amount within the conduits in penetrating arterioles (Fig. 1o; significantly less, p=0.038). However, because there are 1.9-fold more venules than arterioles in this part of the mouse brain^35^ the total efflux of tau via arterioles and venules was not significantly different (Fig. 1o; 2.2±18% larger in venule conduits, p=0.91), which differs from the prediction of the glymphatic hypothesis that efflux should be solely around venules^9^. These data suggest that tau (and Aβ, see below) exits the brain parenchyma in the conduits in penetrating arteriole walls and around ascending venules, enters the pial-arachnoid mesh of conduits, and thus exits the brain either to dural lymphatics or via conduits in the cranial nerves (see below).

### The exit routes observed operate without pressure elevation

Fluorescently labelled tau-441 (1 µM), applied without pressure^36^ by topical application to the CSF space for 5 minutes through a cranial window following removal of the dura in 6-10 week-old mice (n=3), rapidly distributed (within 5-10 mins) within the brain parenchyma and could be seen more brightly around cortical capillaries (Extended Data Fig. 3a), within the smooth muscle wall of penetrating arterioles and around ascending venules (Extended Data Fig. 3b). From these conduits, the tracer dispersed throughout the pial and inner arachnoid layers, again displaying a reticular, web-like pattern (Fig. 2a-d; Extended Data Fig. 3a-d). The pial arachnoid conduits along which the fluorescent tau tracked were spatially organised, with prominence around pial arteries (Fig. 2a-d; Extended Data Fig. 3c-d), more diffusely along the surfaces of pial veins (Fig. 2c; Extended Data Fig. 3c-d), and at locations where pial arteries and pial veins crossed and their associated conduits merged (Fig. 2d). Tracer could also be observed tracking through spire- or veil-like extensions arching from pial artery walls to pial veins and to the wider pial-arachnoid conduit mesh, and within strand-like structures (spanning the CSF space) connecting the pia to the inner arachnoid layer (Fig. 2a-c).

**Fig. 2.**
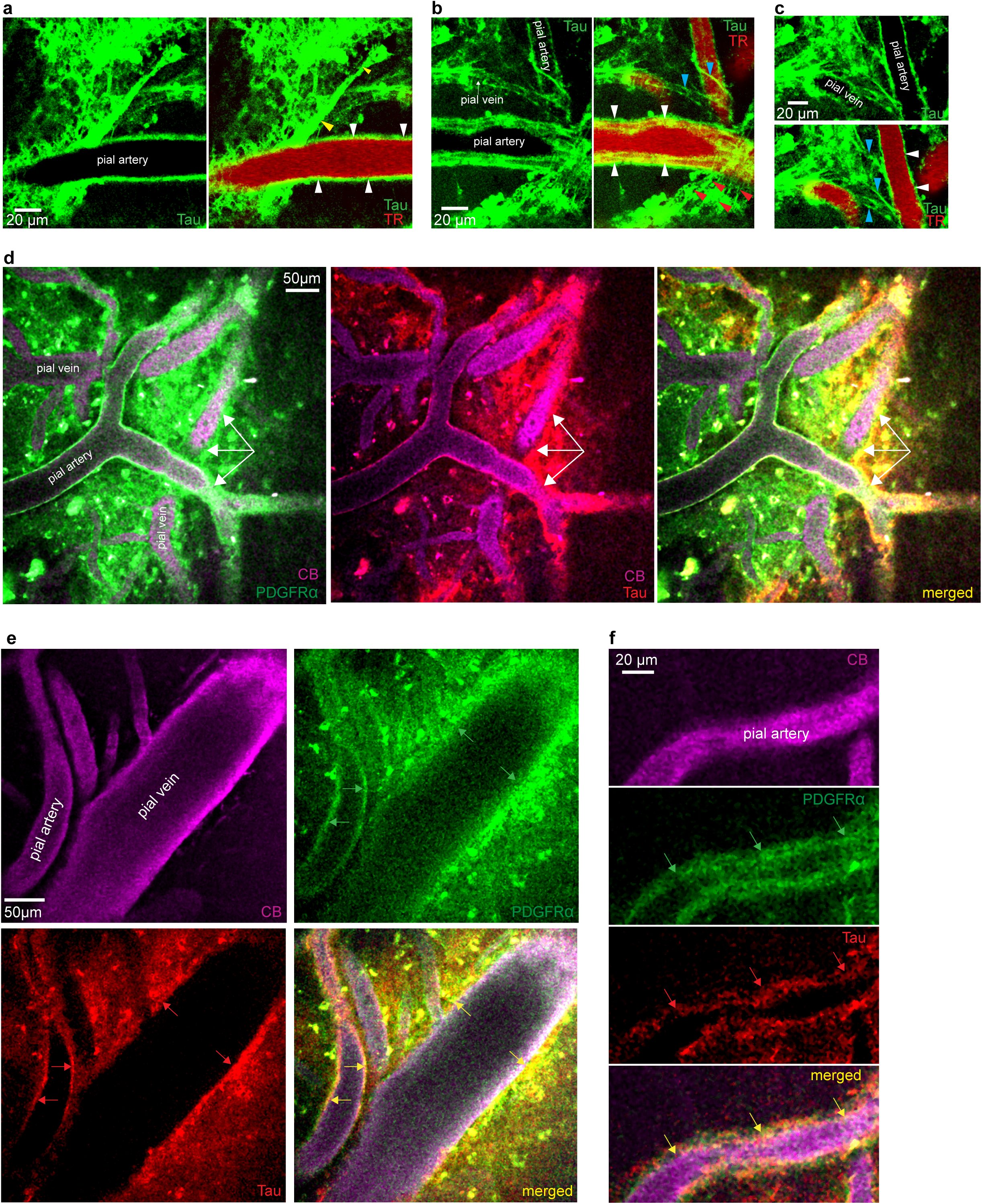
Tau exits the brain via conduits formed by PDGFRα expressing fibroblasts.(a-c) Two-photon imaging at the pial surface after topical tau application (with Texad Red, TR, in blood) reveals that monomeric tau-441 (green) passes from linear conduits (IPAD route) in a pial artery wall (white arrowheads) to the FRC conduit mesh via spire-like (yellow arrowheads, (a)), sail-like (red arrowheads, (b)) and spindle-like (blue arrowheads, (b-c)) connections. **(d)** Similar images 5-min after topical application of fluorescently labelled (red) monomeric tau-441 in 8-week Pdgfra-mGFP-CreERT2 mouse (with green fluorescently-labelled fibroblasts, and with Cascade Blue, CB, magenta, in the blood) showing that drainage occurs via an FRC conduit system (and without the pressure elevations inherent to intra-parenchymal injections of tracer). White arrows highlight tau coursing through FRC conduits bridging from a pial artery to a pial vein. **(e-f)** Representative in vivo 2-photon images (via cranial window) of 8-week Pdgfra-mGFP-CreERT2 mouse (with green fluorescently-labelled fibroblasts, green arrows) 5-min post-parenchymal injection of fluorescently-labelled (red) monomeric tau-441 (red arrows), with Cascade Blue (CB, magenta) in blood. The clearance of tau within penetrating arteriole and pial artery walls, around ascending venules and pial veins, and throughout the pial arachnoid layers occurs via conduits formed by fibroblasts reticular cells (FRC, yellow arrows).

### Exit of tau occurs via fibroblast reticular cell (FRC) conduits

Some experiments using intracortical micro-injection or topical tracer application via a cranial window were performed in Pdgfra-mGFP-CreERT2 mice^37,38^ (which have green fluorescently labelled fibroblasts). This revealed that the conduit network within the penetrating arteriolar and pial artery walls, surrounding the ascending venules and spanning the pia and inner arachnoid layers, through which tau (and Aβ: see below) coursed, labelled for PDGFRα, suggesting^16^ that it was composed of fibroblast reticular cells (FRCs), with a denser FRC conduit mesh coverage of pial veins than of pial arteries (Fig. 2d-f, tau tagged with a red fluorophore was used for these experiments).

To ensure that these findings did not reflect a confounding effect of removing a small portion of skull and dura, we injected fluorescently labelled tau-441 (5 µl over 5 min, a lower rate than used by others^9,11,14,39^) into the mouse cisterna magna, from where it passes to the cerebrospinal fluid bathing the brain (without creating a cranial window). Mice were perfusion-fixed after 5-10 minutes, 1 hour and 3 hours (n=3 for each), and brain slices obtained from the tissue were antibody-labelled before confocal imaging. This revealed that the PDGFRα-labelled conduits in the pial-arachnoid layers also label for the lymphatic markers podoplanin and Prox1 (Fig. 3a). At 5–10 minutes, tau was only visible enveloping the outer surface of penetrating arterioles and pial arteries (and not present in the surrounding pial-arachnoid FRC conduit mesh), capturing the phase of tracer inflow from the CSF into the parenchyma along the subarachnoid paravascular space (as previously described^9,10,29^; Fig. 3b). At later times, fluorescent tau was localised within the smooth muscle layer of penetrating cortical arterioles and pial arteries (as above for parenchymal injection, and as reported previously^29^) and also seen within FRC conduits (displaying a mesh-like, reticulated morphology) in the nearby pial-arachnoid layers (Fig. 3c-e). Fluorescently labelled tau was only observed around ascending venules and deep cerebral veins between one and three hours after intracisternal magna (ICM) injection (Fig. 3f, Extended Data Fig. 4a), consistent with earlier reports^9^. These clearance routes are as seen for intra-parenchymal injection, except that the tracer takes longer to reach the ascending venules, presumably because it first has to enter the extracellular space of the parenchyma from the Virchow-Robin space around arterioles and then diffuse to the venules.

**Fig. 3.**
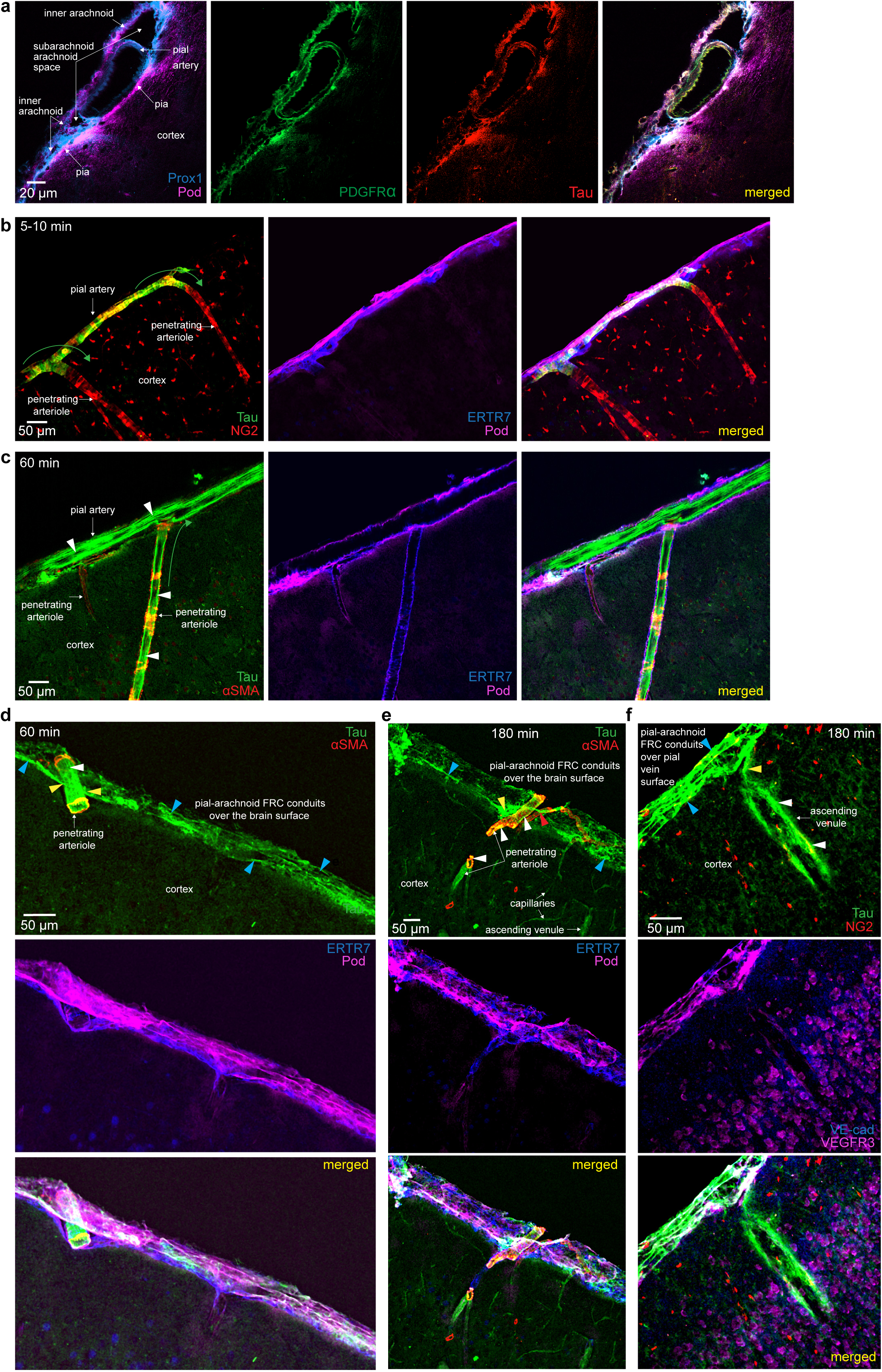
Exit of tau occurs via fibroblast reticular cell (FRC) conduits. **(a)** Ventral surface of tissue-cleared brain 60-min post-ICM injection reveals tau-441 in PDGFRa-labelled conduits in pial artery walls, that connect to the pial/arachnoid conduit mesh via conduits labelling for the lymphatic markers podoplanin (Pod, magenta), Prox1 (blue) and PDGFRα. **(b-f)** Confocal images of fluorescently labelled (green) tau-441 following ICM injection and perfusion-fixation. **(b)** 5-min post-ICM injection, tau is in the periarterial subarachnoid space and entering (green arrows) the Virchow-Robin space (as found by others^9,29,47^) with no tau in the cortical parenchyma. Pial-arachnoid conduits label for collagen VI (ERTR7, blue) and podoplanin (Pod, magenta). NG2 labels smooth muscle cells and pericytes (and oligodendrocyte precursor cells). **(c)** 60-min post-ICM injection, tau is present dimly throughout the cortex and draining (green arrow) via linear channels within penetrating arterioles and pial artery walls (labelled for smooth muscle actin, α-SMA, red), as well as in the surrounding pial conduits (white arrowheads) labelling for collagen VI (ERTR7, blue) and podoplanin (Pod, magenta). **(d)** 60-min post ICM-injection, tau is seen again dimly within the cortex, linear channels (white arrowheads) of a penetrating artery wall (α-SMA, red)) and throughout FRC conduits (blue arrowheads) coursing through the pial-arachnoid layers labelling for collagen VI (ERTR7, blue) and podoplanin (Pod, magenta). Note the spire-like connections (yellow arrowheads and as seen in vivo, Fig. 2a) along which tau-441 passes from the pial artery wall to the pial-arachnoid FRC conduit mesh. **(e)** 180-min post-ICM injection shows tau in the cortex, surrounding cortical capillary-sized vessels (blue arrowheads) and within the wall (white arrowheads) of a penetrating arteriole (becoming a pial artery where it reaches the surface) with connection to the pial-arachnoid FRC mesh (labelling for collagen VI (ERTR7, blue) and podoplanin (Pod, magenta) via both a sail-like (yellow arrowhead) and a spire-like (red arrow) fibroblast structure (as also seen in vivo, Fig. 2a-b). **(f)** 180-min post ICM injection, reveals clearance of tau to the pial-arachnoid FRC conduit mesh (labelling for VE-cadherin, VE, blue; VEGFR3, magenta) covering a pial vein lacking NG2 expression (blue arrowheads) via channels along the outer aspect of an ascending venule (white arrowheads).

ICM injection of a much smaller volume of fluorescently labelled tau (500 nl over 5 minutes) with 4% perfusion fixation performed after 90 minutes also showed tracer dispersed within the pial arterial walls, and in conduits around venules and in the pial-arachnoid FRC mesh (n=3, Extended Data Fig. 4a-c).

### Additional markers of the conduit system

In the penetrating arteriole and pial arterial walls, the PDGFRα-labelled conduits along which the applied fluorescent molecules tracked out of the brain labelled for VE-cadherin (Fig. 4a) and the pial sheath around the penetrating arterioles labelled for the lymphatic conduit markers^17^ podoplanin and ERTR7 (Fig. 3c-e; podoplanin labelling was stronger around the pial than around the penetrating arterial conduits). Over the brain surface, conduits in the pial arteries connect with, and transfer tau to, the wider FRC conduit mesh within the pia and inner arachnoid membranes covering the brain surface and pial veins, which label for podoplanin (Fig. 3a-e), VEGFR3 (Fig. 3f, Fig. 4b), and (mainly in the inner arachnoid) Prox1 (Fig. 3a) and CRABP2 (Extended Data Fig. 4e). Similarly, fluorescently labelled tau tracked from podoplanin and ERTR7 labelled conduits around ascending venules (Ext Data Fig 4a) into VE-cadherin and VEGFR3 labelled conduits in the pia-arachnoid (Fig. 3f).

**Fig. 4.**
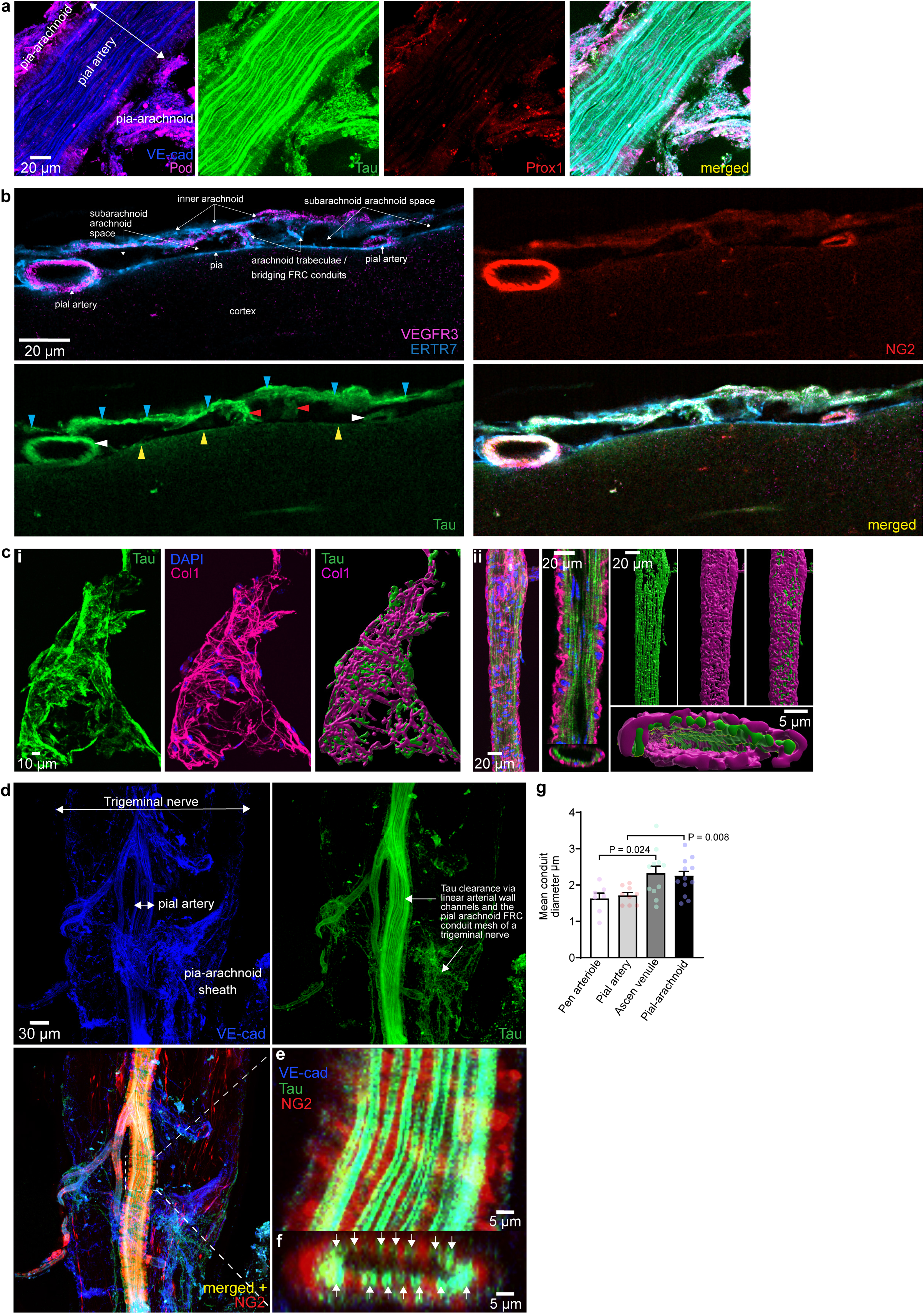
Microstructure of FRC conduits in the pia-arachnoid and exiting cranial nerves.(a) In the pial layer 60 min post-ICM injection, fluorescently labelled monomeric tau-441 (green) clears through VE-cadherin (VE-cad, blue) labelled longitudinal conduits running along a pial artery wall and within surrounding pial-arachnoid FRC conduits labelling for VE-cadherin, Prox1 (red) and podoplanin (Pod, magenta)). (**b**) Tau is distributed within the walls (white arrowheads in tau panel) of pial arteries (NG2, red) and throughout the pia (yellow arrowheads) and inner arachnoid layers (blue arrow heads), which are connected by strand-like fibroblast and ECM component extensions (red arrowheads) labelled for collagen VI (ERTR7, blue) and VEGFR3 (magenta). **(c)** 60-mins post-ICM injection, fluorescently labelled tau-441 (**i**) closely associates with collagen I fibres (Col1, magenta; DAPI, blue) within the pial and arachnoid layers surrounding a large vein, and **(ii)** locates to longitudinally orientated channels enveloped by collagen I fibres within an artery wall (highlighted by 3D rendered images). (**d)** In a trigeminal nerve, 60-min post-ICM injection (in an NG2-DsRed mouse with red pericytes and arteriolar smooth muscle cells), fluorescently labelled tau is cleared via VE-cadherin labelled channels within a nerve pial artery wall and in the wider surrounding pial-arachnoid FRC conduit mesh. (**e**) Enlarged view of boxed area in (d). (**f**) End view of artery in (e) shows green tau-filled conduits around the inner side of the artery wall. (**g**) Width of the tau-filled conduits in 25 mice, in penetrating arterioles, pial arteries, ascending venules and in the pial-arachnoid mesh of conduits.

Several collagen isoforms are associated with the conduit system. The pial arterial wall conduits expressed collagen IV (Extended Data Fig. 4f), and are surrounded by a rim of collagen I (Fig. 4c(ii)). The pial conduits express collagen types I, III and VI (Fig. 4c(i), Extended Data Fig. 4g, Extended Data Fig. 4h) which are part of the FRC structure^20^ (Extended Data Fig. 1f), in keeping with known high expression levels of ECM genes Col1A1, Col3A1 and Col6A1 and Col 6A2 by pial and inner arachnoid fibroblasts^25,40,41^.

Tau tracking through conduits closely associated with collagen VI (ERTR7: Fig. 3c-e; Extended Data Fig 4a, d, h) and collagen I (Fig. 4c) fibre bundles in the FRC mesh is similar to that described for lymph and solute transport within the FRC conduits in lymph nodes^16,20^. The diversity of labelling for different lymphatic and fibroblast markers may reflect heterogeneity of the population of fibroblasts that form these conduits^42^. A summary of the labelling of the conduits in different locations is given in Table 1.

### Conduits bridging the pia and arachnoid layers

Proteins exiting the brain may need to cross from the pial surface to the arachnoid in order to ultimately exit to dural lymph vessels^43^. Conduits labelling for ERTR7 and VEGFR3 were observed to bridge the pial and arachnoid layers at focal locations away from both pial arteries and veins (Fig. 4b), allowing clearance of tau across the subarachnoid space independently of bridging conduits at artery-vein crossover points.

### Conduits within arteries and the pia-arachnoid of exiting cranial nerves

Part of the lymphatic efflux from the brain is thought to be along exiting cranial nerves (Extended Data Fig. 1), so confocal microscopy was applied to optic and trigeminal nerves carefully dissected from the ventral brain surface of mice (post-perfusion fixation), following ICM application of fluorescent tau. The conduits carrying tau out along arteries within the nerves exhibited a regular disposition around the periphery of the artery SMC layer (Fig. 4d-f), and showed a clear “train track” like morphology, perhaps reflecting tau moving along the extracellular space of FRC conduits either side of a central collagen core (Extended Data Fig. 1f), whilst conduits in the pial-arachnoid layer around the nerve again possessed a mesh-like morphology (Fig. 4d).

### Dimensions of FRC conduits

Measuring the width of the conduits from the breadth of the tau-filled channels showed that they spanned a range of diameters, with a mean value averaged over all locations of 2.03±0.09 μm, and a clear difference in diameter between the more linear conduits within penetrating arterioles and pial arteries (∼1.7 μm) and the less organised conduits around ascending venules and in the pial-arachnoid layer (∼2.3 μm, significantly different, p=0.024 comparing penetrating arterioles and ascending venules, and p=0.008 comparing pial arteries and the pial-arachnoid mesh, Fig. 4g).

### Aβ_1-42_ exits via the same conduits as tau

After parenchymal injection of fluorescently tagged amyloid β_1-42_ (as performed above for tau), Aβ was observed to track along conduits in the wall of penetrating arterioles and pial arteries and around ascending venules (Fig. 5a-b), to enter conduits spreading extensively though the pial-arachnoid layers (Fig. 5c-d), and to leave the brain via the pial-arachnoid conduit mesh around the optic and trigeminal nerves (expressing VEGFR3, VE-cadherin and Prox1: Fig. 5e-f), and through linear conduits within the arteries of these nerves that express VE-cadherin but not Prox1 or VEGFR3 (Fig. 5f). After ICM injection of Aβ_1-42_ its exit conduits were seen to label for VE-cadherin, and in the pia podoplanin (Fig. 5g-h), as well as ERTR7 and VEGFR3 (Fig. 5i).

**Fig. 5.**
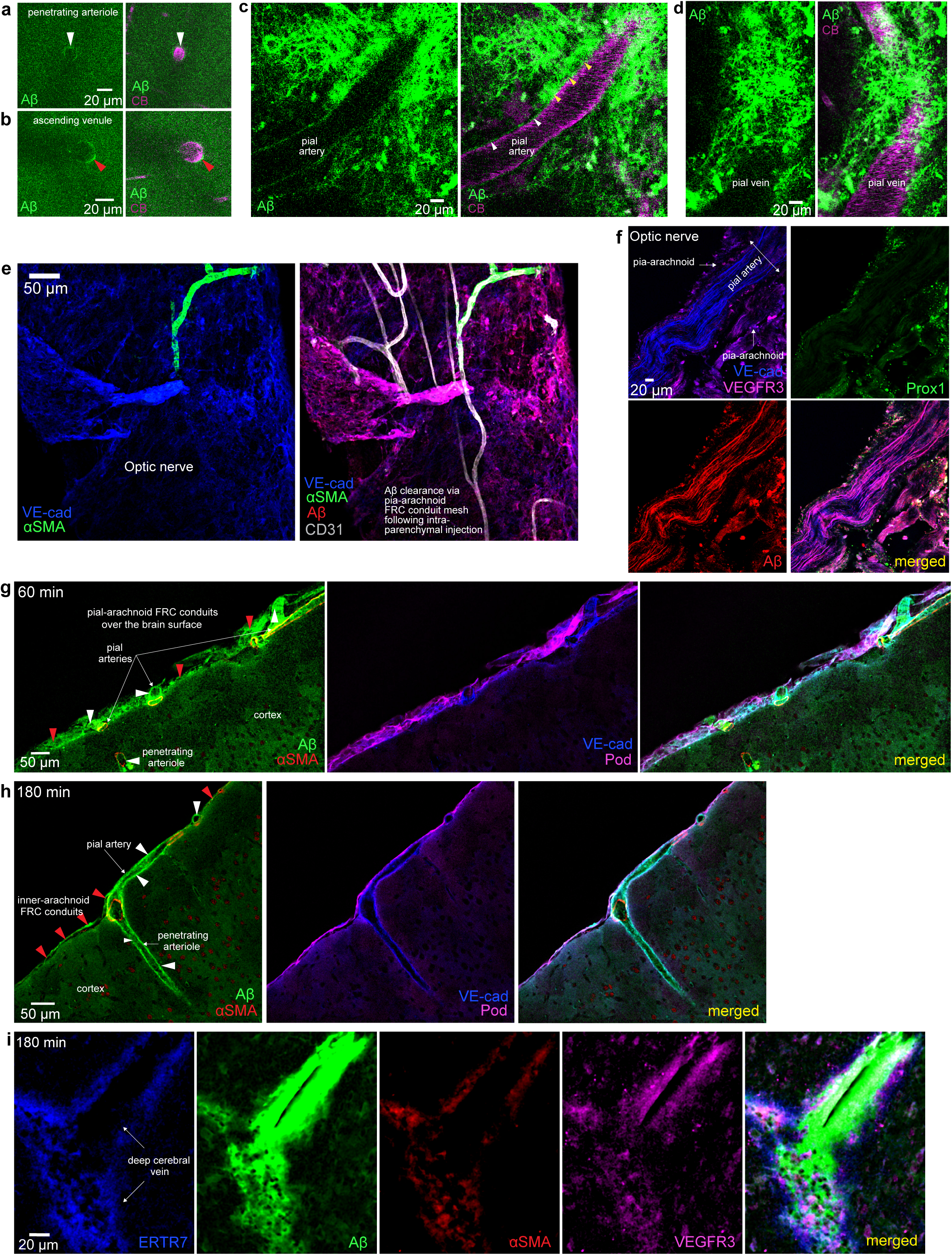
Aβ exits via the same fibroblast reticular cell (FRC) conduits as for tau. (**a-e)** In vivo two-photon microscopy of 8-week-old WT mouse following intra-parenchymal application (at a depth of 250 μm) of fluorescently-labelled (green) Aβ_1-42_ via cranial window (CB = Cascade Blue in blood), showing Aβ in conduits in the wall of penetrating arterioles (**a**), around ascending venules (**b**), in the walls of a pial artery (**c**), in the pial-arachnoid mesh around a pial vein (**d**), in pial-arachnoid web-like conduits expressing VE-cadherin surrounding an optic nerve (**e**) and in linear conduits expressing VE-cadherin (but not VEGFR3 or Prox1) running along an artery in the optic nerve (**f**). **(g-i)** 180-min post ICM-injection, Aβ_1-42_ is being cleared via VE-cadherin expressing channels (white arrowheads) in the pia (**g**) and in an α-SMA (red) expressing penetrating arteriole and pial artery (which also expresses podoplanin) and throughout the FRC conduits (red arrowheads) coursing through the pial-arachnoid layers (**h**). **(i)** 180-min post-ICM injection shows clearance of Aβ_1-42_ via conduits associated with ERTR7 (collagen VI, blue) and labelling for VEGFR3 (magenta) surrounding a large, deep cerebral vein (only weakly labelled with smooth muscle actin antibody, α-SMA, red).

### The gross organisation of exit conduits

To obtain a macroscopic perspective of the spatial distribution of the FRC conduit network over the brain surface we used light-sheet and confocal microscopy to image larger volumes of tissue-cleared, antibody-labelled brain sections from WT and NG2dsRed mice (Fig. 6a-b, Extended Data Fig. 5a-c). This showed that the FRC mesh enwraps and penetrates the parenchyma of the entire mouse brain, including the dorsal and ventral surfaces, the olfactory bulb, the brainstem, and the cerebellum, as well as the exiting cranial nerves, as described above. The FRC conduit mesh was labelled in some regions with an antibody recognising Lyve-1 (which displayed colocalisation with podoplanin labelling), though it is possible that the antibody recognised CD206-negative perivascular macrophages with unusual morphology (Extended Data Fig. 5d). Non-macrophage LYVE-1 labelling in the leptomeningeal layers has, however, been described previously^44^.

**Fig. 6.**
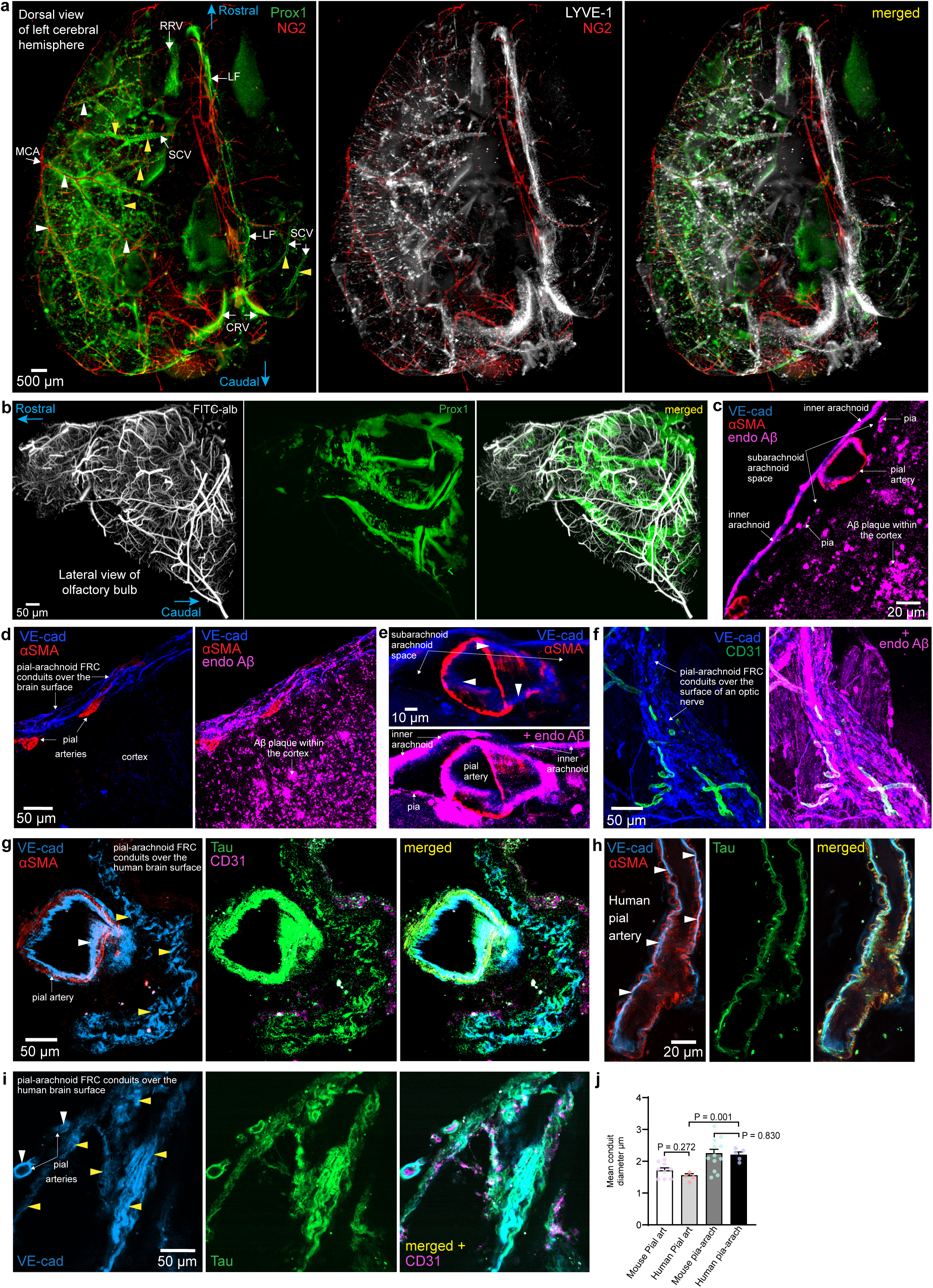
The global FRC conduit network, its clearance of endogenous Aβ and its presence in human brain. **(a-b)** Light sheet images of tissue-cleared cortical hemispheres from 8-week-old NG2DsRed mouse (performed 5 times). **(a)** Dorsal view of the spatial distribution of the FRC conduit mesh of the inner pia-arachnoid, labelling for Prox1 (green) and perivascular macrophages (LYVE-1, white), pial arteries and arterioles expressing NG2 (red, yellow arrowhead) and superior cerebral/pial veins (NG2 –ve, white arrowhead). Prominent labelling is around the regions of the middle cerebral artery (MCA) and its branches, a superior cerebral/pial vein (SCV), the rostral rhinal vein (RRV), the caudal rhinal vein (CRV) and the longitudinal fissure (LF). **(b)** Distribution of the pial-arachnoid FRC conduit labelling for Prox1 (green) around the olfactory bulb. **(c-f)** Confocal images from 9-month APP^NL-G-F^ mice show endogenous Aβ (endoAβ, magenta) deposited within: **(c)** a pial artery wall (labelling for α-SMA (red)) and in the wider pial arachnoid FRC conduits (labelling for VE-cadherin (VE-Cad, blue)); **(d)** the mesh-like pial-arachnoid FRCs of the brain surface; **(e)** VE-cadherin labelled channels (white arrowheads) within a pial artery wall; **(f)** pial-arachnoid FRC conduits labelling for VE-cadherin (blue) surrounding an optic nerve from a 9-month-old APP^NL-G-F^ mouse. **(g-i)** Confocal images of live human cortex incubated for one hour with fluorescently labelled (green) monomeric tau-441. The tau is seen in VE-cadherin-labelled conduits (white arrowhead) within a pial artery wall labelling for smooth-muscle actin (α-SMA, Red), and in mesh-like VE-cadherin expressing (blue) pial-arachnoid conduits (yellow arrowheads).

The distribution of the FRC conduit mesh was not uniform over the brain, displaying prominence around the paravascular region of middle cerebral artery branches, the surface of pial veins and the rostral and caudal rhinal vein regions, as well as the longitudinal and cerebrocerebellar fissures (Fig. 6a; Extended Data Fig. 5b, templating the distribution of dural lymphatic locations). Where pial arteries and veins overlap, FRC conduits within the pia and arachnoid layers fuse, allowing (as described above in vivo) solutes to flow directly from around and within the arterial muscle layer to around pial veins. Further linking pathways allowing egress of CSF/ISF solutes from the periarterial regions to the conduits surrounding veins (and presumably on to the dural lymphatics^31^), are provided by the spire-or veil-like extensions noted above (Fig. 2a-c; Extended Data Fig. 3c) which arch from pial artery walls to pial veins and to the wider pial-arachnoid conduit mesh, and by strand-like structures (spanning the CSF space) connecting the pia to the inner arachnoid layer at focal locations (Extended Data Fig. 5f). On the ventral and dorsal surfaces of the brain, respectively, tracer could be found tracking through FRC conduits within arachnoid trabeculae and more delicate strand-like fibroblast and ECM component extensions, respectively, providing a pathway for solutes to move across the CSF space towards the dura (Fig. 4b, Extended Data Fig. 5f).

### Endogenous Aβ is cleared by the same pathway as exogenous tau and Aβ

We examined the clearance pathway of endogenously expressed mutant human Aβ in 9-month-old APP^NL-G-F^ Alzheimer model mice^34^, using confocal imaging of antibody-labelled fixed tissue. This revealed the accumulation of human mutant Aβ in the VE-cadherin expressing FRC conduit mesh within the pial-arachnoid layers surrounding both pial arteries and pial veins, as well as within the pial arterial muscle wall (Fig. 6c-e). The topographical distribution of accumulated endogenous Aβ in the pial FRC conduit mesh of 9-month-old mice was remarkably similar to that observed following application of fluorescent tracer (tau and Aβ) via intracortical or ICM injection (Extended Data Fig. 5g), or through a cranial window. Examining the optic and trigeminal nerves of these 9-month-old APP^NL-G-F^ mice showed heavy accumulation of endogenous Aβ along the VE-cadherin labelled channels of the enveloping pial-arachnoid FRC conduit mesh (Fig. 6f).

### Similar conduits exist in human brain

We investigated whether the same FRC conduits exist in the human brain, using cortical living tissue that would otherwise be discarded during neurosurgery on patients with glioma. Monomeric tau-441 (and Aβ_1-42_, Extended Data Fig. 5h), applied for an hour in solution bathing brain slices made from the excised tissue, both located selectively to FRC conduits within artery walls and within the pial-arachnoid layers (including around pial veins) that labelled for VE-cadherin (Fig. 6g-i), podoplanin, collagen I and PDGFRα (Extended Data Fig. 5h-i) rather than occupying the cortical parenchyma, suggesting the presence of active uptake into the conduits, and implying a conservation of the conduit removal mechanism between mice and humans. The dimensions of the conduits in human cortex samples from 5 people were similar to those in mice (Fig. 6j), with a mean diameter across all conduits of 1.94±0.07 μm.

## Discussion

We have shown that, in mice, fluorescently labelled neurodegeneration-inducing molecules injected into the brain parenchyma, injected into the CSF in the cisterna magna or applied topically, rapidly drain into a previously undefined web-like spatially-organised conduit system, which routes molecules up the wall of penetrating arterioles (continuing as pial arteries) and up around ascending venules, from where they move into a pial-arachnoid conduit network enveloping the brain surface, pial veins and cranial nerves. From here molecules may efflux via bridging veins and arachnoid cuff exit points^31^ to the dural lymphatics, or via the cribriform plate or along cranial nerves^10–14^. Present also around the human brain, this drainage mesh is formed of reticular fibroblasts labelling for VE-cadherin^15^ and the classic lymphatic markers^17^ podoplanin, Prox1, VEGFR3 and their ECM components collagen fibres I, III, IV and VI. A similar fluid- and solute-transporting conduit system formed of reticular fibroblasts expressing podoplanin has been well characterised in lymph nodes and secondary lymphoid organs^16,20^, and those conduits have a diameter (median value^45^ 2.5 μm) similar to that which we measured in Fig. 4g.

Previous controversy on how neurotoxic molecules are lost from the brain has focussed on efflux via intramural periarterial drainage (IPAD, within the wall of penetrating arterioles) versus efflux around ascending venules driven by water influx through aquaporin 4 expressing astrocyte endfeet apposed to penetrating arterioles (the glymphatic system). It turns out that aspects of both of those proposed systems are correct in that parenchymal tau and Aβ do move to the pial surface in conduits both within the wall of penetrating arterioles and surrounding ascending venules. However, our quantification of efflux shows an approximately equal molecular flux within arteriole walls and around venules, while IPAD posits efflux to be largely via arteriole walls and the glymphatic idea proposes efflux solely around venules. Given the cellular similarities between the conduits within arteriole walls and around venule walls it is economical to hypothesise that the same mechanism operates in both locations (Extended Data Fig. 1e), and is mediated by conduits of the form already known to exist in lymph nodes (Extended Data Fig. 1f). Based on our knowledge of the homologous conduits in lymph nodes, it is likely that the brain conduit system imposes a molecular weight limit to the size of molecules removed^20^ of the order of 70 kD. This would allow the removal of a wide range of soluble Aβ oligomers (monomeric MW 4.5 kD), but only the removal of tau monomers (MW 46-70 kD).

Defining the cellular and molecular composition of these pathways will facilitate investigation of questions about their function. The higher fluorescence of tagged tau and Aβ within the conduits than in the surrounding parenchyma strongly suggests an accumulation mechanism, the molecular basis of which is unknown (although an intra-conduit environment that potentiates fluorescence cannot be ruled out). Tracer can rapidly travel large distances along conduits, favouring solute movement by bulk flow over diffusion. How fluid and solutes are pumped with such speed is unknown (reaching FRC conduits within the pial and arachnoid layers and the arterial walls of vessels on the dorsal surface of the mouse brain within 7.5 minutes post-parenchymal or ICM injection). Rhythmic compression by blood vessel pulsation^8^ or active pumping in the dural vessels^46^ or in conduits in the exiting cranial nerves may contribute. It is also unknown whether fluid and solute flow through the FRC conduit meshwork can be modulated by physical, physiological, or pharmacological means; for example, there is controversy over how molecule clearance is altered during sleep^47,48^ and by alteration of astrocyte aquaporin expression^9^.

Despite the advance inherent in defining these conduits, our work comes with some cautions. Although manipulation of more distal lymphatic components of outflow can both alter clearance of molecules from the brain^49–51^, an unquantified fraction of endogenously generated neurotoxic molecules may be degraded within the brain or leave across vascular endothelial cells to the blood. We assume that parenchymally-generated Aβ and tau would enter the conduits within the parenchyma and be cleared towards the pia (as we observe for endogenous Aβ in an AD model mouse in Fig. 6c-f), but we cannot exclude the possibility that a fraction of neurotoxic solutes may enter the FRC conduit mesh within the pial-arachnoid layers directly from the subarachnoid space and/or periarterial space (without first entering the brain)^52^ or from the smooth muscle basement membrane of arterioles^29^, nor that a portion of fluids and solutes leave the FRC conduits to enter the subarachnoid space. A point of controversy has been whether injecting substances into the brain parenchyma or cerebrospinal fluid alters the exit pathways followed by neurotoxic molecules. We found that tracer was similarly distributed throughout the FRC conduit system in mice following intra-parenchymal injection, low-volume (0.1 μl/min for 5 min) ICM injection of tracer into the CSF, or application of tracer to the CSF via a cranial window without pressure applied, or following incubation of live human brain tissue slices with fluorescent Aβ and tau. One could argue that all of these methods are artificial situations, so the most convincing demonstration that our data are relevant to normal physiology and pathophysiology is that our identification of the conduits involved has allowed us to demonstrate that they are employed for the removal of endogenous Aβ in Alzheimer’s model mice (Fig. 6c-f).

The distribution of Aβ deposits along the FRC conduits in APP^NL-G-F^ mice (Fig. 6c-f) was morphologically similar to that of tracer localising throughout the FRC conduits following its injection into the parenchyma or cisterna magna, i.e. in a pathway from parenchymal arterioles and ascending venules, to pial arteries and pial veins, and to exiting cranial nerves. The presence of Aβ in the conduits around pial veins (branches of bridging veins) *in vivo* and in fixed tissue obtained from 9-month-old APP^NL-G-F^ mice (Extended Data Fig. 5g) indicates that this may be a vital efflux route for waste solutes to the dural lymphatics via recently described ACE points^31^, or to conduits in exiting cranial nerves (Fig. 4d-f). Marked amyloidosis of bridging cerebral veins and the dural regions surrounding the points at which bridging veins cross into the dura (seen in both Alzheimer’s disease model mice and humans), suggests that the FRC conduits within the pial-arachnoid layers may deliver fluid and waste to the dura via ACE points^43^. Any slowing or inadequacy, in the face of increased Aβ production, of this exit route might be expected to lead to Aβ accumulation earlier in the pathway, for example in the walls of arterioles, as is seen in the pathology of amyloid angiopathy.

Drainage of tracer through the FRC conduit mesh surrounding the optic nerve and trigeminal nerves (Figs. 4d, 5e-f) is in keeping with previous findings that CSF and solutes drain out of the brain via a perineural route (see Introduction). Whether the FRC conduits in the pial-subarachnoid layers deliver fluid and solutes to dural lymphatics located around exiting cranial nerves^4,10,13,53^, or via gaps in the ABCL, or drainage continues along a true perineural route to reach extracranial lymphatics, requires further investigation. Nevertheless, our data unify previous observations of brain clearance of solutes from the CSF/ISF within penetrating arterioles/pial arterial walls^8,28,29,54,55^, the paravascular region of ascending venules, pial and deep cerebral veins^9,31,43,55^ and cranial nerves^4,10,13,53^, into a single working model (Extended Data Fig. 1e) and can explain how fluids and solutes can cross the subarachnoid space from the brain to the inner arachnoid via FRC conduits.

The definition of a brain drainage conduit system has implications for understanding the aetiology of diseases in which blockage, degeneration or injury to this delicate mesh (from recurrent collisions during contact sports^56–58^, for example, or CAA^59^) impairs the clearance of fluid and solutes from the brain and spinal cord parenchyma, potentially leading to oedema and the accumulation of toxic metabolites, microbial antigens, and neurotoxic proteins such as Aβ, p-tau, α-synuclein and TDP-43. Disruption of the FRC conduit system may also impair immune surveillance and the response to antigens that are ordinarily presented as they drain from the central nervous system.

## Methods

### Animals

Mice (of either sex) in individually ventilated cages (12-h light/dark cycle, 21±1 °C, 45– 65% humidity) were fed normal chow and tap water ad libitum. Cages contained Aspen chip and Sizzle-Nest bedding and an environmental enrichment tunnel. We used the following transgenic mouse lines, in addition to wild type (WT) mice, for our experiments: NG2-dsRed mice^60^ (expressing a red fluorescent protein (DsRed.T1) in cells that produce nerve-glial antigen 2 (NG2), driven by the *Cspg4* promoter/enhancer), Pdgfra-mGFP-CreERT2^37,38^ and APP^NL–G–F^ (Alzheimer’s disease model^34^) mice (which show increased Aβ production and aggregation (via Swedish, Arctic and Iberian mutations of APP), with mutated APP being knocked-in at its endogenous locus to avoid overexpression artifacts). All mice were euthanised by cervical dislocation or cardiac perfusion under terminal anaesthesia. Animal breeding, experimental procedures and euthanasia methods were in accordance with UK Home Office regulations (Guidance on the Operation of Animals, Scientific Procedures Act, 1986) and advice from the UCL Animal Welfare Ethical Review Board.

### Experiments in vivo

Mice aged 6-10 weeks were anaesthetised with urethane (1.55 g/kg, given in two doses 15 min apart). Adequate anaesthesia was ensured by confirming the absence of a blink reflex and withdrawal response to a paw pinch. Body temperature was maintained at 36°C, and eyes were protected from drying by applying polyacrylic acid eye drops (Dr. Winzer Pharma).

### In vivo 2-photon imaging with tracer applied via cranial window or intracortical injection

Mice were secured in a stereotaxic frame, and lidocaine/prilocaine (AstraZeneca) was applied topically before exposing the skull. A custom-built headplate was then attached to the skull using superglue to create a sealed well to contain HEPES-buffered aCSF (140 mM NaCl, 10 mM HEPES, 2.5 mM KCl, 1 mM NaH_2_PO_4_, 10 mM glucose, 2 mM CaCl_2_, 1 mM MgCl_2_, pH 7.4) during imaging. A craniotomy of approximately 3 mm diameter was performed over the right primary somatosensory cortex, and the underlying dura was carefully resected. During imaging, the headplate was secured under the objective on a custom-built stage.

Texas Red (2%) or Cascade Blue (1%) were injected retro-orbitally or via the tail vein to delineate the brain vasculature. Two-photon excitation employed a Newport-Spectraphysics Ti: sapphire MaiTai laser pulsing at 80 MHz, and a Zeiss LSM710 microscope with a 20× water immersion objective (NA 1.0). Fluorescence was evoked using a wavelength of: 920 nm for DsRed, Texas Red and Hylite 555-labelled monomeric tau and Aβ_1-42_; 820 nm for Hylite 488-labelled monomeric tau and Aβ_1-42_; and 800 nm for Cascade-Blue. The mean laser power under the objective did not exceed 35 mW. Pial arteries and pial veins, and their respective branches - penetrating arterioles and ascending venules - could be identified by morphology, branching pattern, and by the typical ring shape of vascular smooth muscle cells in pial arteries and penetrating arterioles expressing DsRed in NG2-DsRed mice. Capillaries could be identified according to size (<10µm), branch order relative to penetrating arterioles and ascending venules and the presence of enwrapping pericytes expressing DsRed in NG2-DsRed mice. Image stacks were taken in 1 μm depth increments across layers I to IV of the cortex (up to 200 μm deep from the cortical surface).

Monomeric fluorescent tau (human recombinant, HiLyte Fluor 488- or 555-labelled Anaspec Tau-441, AS-56085 or custom order) or monomeric Aβ_1-42_ (Anaspec HiLyte Fluor 488- or 555-labelled β-Amyloid_1-42_, AS-60479-01 or AS-60480-01, reconstituted by adding 50 µl 1% NH_4_OH to 0.1 mg β-Amyloid_1-42_ and diluted to 1 mg/ml with PBS) were employed. They were applied, either to the surface of the cortex via a cranial window in 150 µl in HEPES-buffered aCSF (as for human slices below) containing 1 μM tau-441 or 44 µM Aβ_1-42_ for 5 minutes before removal (and surface wash with aCSF), or were injected into the cortex (500 nl) at a rate of 100 nl per minute using a 34G needle (inserted through a small hole drilled into the skull lateral to the cranial window, so that the needle tip was located deep to the area being imaged) connected to a pump (UMP3 UltraMicroPump III) at concentrations of 10 μM for tau-441 or 89 μM for Aβ_1-42_ in HEPES-buffered aCSF.

### In vivo intra-cisterna magna (ICM) injection

Mice were anaesthetised (as per in vivo 2-photon imaging experiments) and their neck shaved and cleaned with a solution of iodine in 70% (v/v) ethanol before being placed in a stereotactic frame (World Precision Instruments). The skin over the back of the neck and the outermost nuchal muscles were both cut, and the inner layer retracted to expose the cisterna magna. A 5 μl Hamilton syringe with a 36-gauge (World Precision Instruments Bevelled NanoFil) needle was used to inject 0.5 or 5 μl of tracer at a rate of 0.1 or 1 μl/minute respectively. Identical results were observed with the different infusion rates. The needle was kept in place for an additional 5 minutes after the injection was complete before being carefully removed. Mice recovered on a heating pad to maintain body temperature at 36°C before cardiac perfusion with 4% PFA in PBS after 5-10 minutes, 30 minutes, 60 minutes or 3 hours. Tracers used were: 2.5 µM monomeric fluorescent tau-441 or monomeric 89 µM Aβ_1-42_ (as described above).

### Mouse brain tissue slicing and optic and trigeminal nerve preparation

Anesthetized mice (1.55 g kg^−1^ urethane) were cardiac perfused with 20ml of warm (34-37°C) PBS followed by ice-cold PFA or with 20 ml of warm (34–37 °C) PBS with heparin (20 IU), followed by 20 ml of warm 0.25% (w/v) FITC-albumin (Sigma-Aldrich, A9771) in 5% (w/v) gelatin from porcine skin (Sigma-Aldrich, G1890) in PBS. FITC-perfused mice were placed head down into ice for 30 min. Brains were extracted and the optic and trigeminal nerves carefully dissected from the ventral surface before being drop fixed in 4% PFA overnight. and 100-μm-thick slices were cut in PBS using a Leica VT1200S vibratome. Following cardiac perfusion with 4% PFA, brains from mice post 2-photon in vivo imaging and ICM tracer injection or brains from mice without preceding experimentation were removed from the cranium, and the optic and trigeminal nerves were carefully dissected from the ventral aspect of the brain. The brains, optic nerves, and trigeminal nerves were then transferred to 4% PFA in PBS at 4 °C for 24 hours and thereafter washed 3 times for 10 mins in PBS. Brains were then either sliced into 100 µm sections in PBS or (as for optic nerves and trigeminal nerves) kept whole and prepared for immunocytochemistry with or without tissue clearing.

### Human brain tissue slices

The work on fresh human brain tissue received ethical approval from the National Health Service (REC number 15/NW/0568). Live human cortical tissue was from 5 patients (2 male and 3 female) undergoing neurosurgical glioma resection at the National Hospital for Neurology and Neurosurgery, Queen Square, London, and informed consent was obtained from all patients (who were not financially compensated). During their neurosurgical operations, apparently normal cortical tissue that was removed (to gain access to the tumour), which would otherwise have been discarded, was transported to the laboratory in HEPES-buffered aCSF at 36°C containing 140 mM NaCl, 10 mM HEPES, 2.5 mM KCl, 1 mM NaH_2_PO_4_, 10 mM glucose, 2 mM CaCl_2_, 1 mM MgCl_2_, pH 7.4 (oxygenated by gassing with 100% medical O_2_) with or without monomeric fluorescent tau (1 µM) or Aβ_1-42_ (2.2 µM) (as described above). After 45 minutes of incubation, the tissue was fixed in 4% paraformaldehyde (PFA) in PBS at 4°C for 24 hours, washed three times in PBS and then sliced into 150-200 µm sections using a Leica VT1000S vibratome. Each patient’s tissue typically generated ∼2 brain slices.

### Immunohistochemistry of human or mouse brain slices and optic and trigeminal nerves

Following incubation in blocking buffer (3% bovine serum albumin/0.5% Triton X-100 in PBS) at room temperature for 1 hour, brain slices and optic and trigeminal nerves were incubated in blocking buffer overnight at 4°C in with the following primary antibodies: anti-Prox1, 1:100 dilution, Angiobio 11-002P; anti-podoplanin, 1:100, eBioscience 15257417; anti-VEGFR3, 1:100, R&D Systems AF743; anti-Lyve-1, 1:200, eBioscience 15286957; anti Lyve-1, 1:200, Invitrogen PA1-16635; anti-αSMA, 1:500, Abcam ab5694; anti-CD144 (VE-Cadherin), 1:200, eBioscience 14-1441-81; anti-CD206, 1:200, Fisher Scientific 18704-1-AP; anti-collagen I (COL1A1), 1:250, Abcam ab21286; anti-Collagen III, 1:250, Abcam ab7778; anti-collagen IV, 1:250, Abcam ab19808; anti-ER-TR7 Col VI), 1:500, R&D Systems NB100-64932; anti-CD31 (PECAM-1) 1:200 eBioscience,15296897; anti-β-Amyloid_1-16_ Alexa Fluor® 647, 1:250, BioLegend 803020; anti-CRABP2, 1:500, Fisher Scientific 10225-1-AP.

Then, after washing 3 times for 10 minutes in PBS, brain slices and optic and trigeminal nerves were incubated in blocking buffer overnight at 4°C with the following secondary antibodies at a dilution of 1:500: anti-rat IgG Alexa Fluor 405, Thermo Fisher Scientific A48268; anti-mouse Alexa Fluor 405, Thermo Fisher Scientific A-31553; anti-mouse Alexa Fluor 488, Thermo Fisher Scientific A-21202; anti-mouse Alexa Fluor 633, Thermo Fisher Scientific A-21050; anti-rat Alexa Fluor 633, Thermo Fisher Scientific A-21094; anti-rabbit Alexa Fluor 488, Thermo Fisher Scientific A21206; anti-goat Alexa Fluor 488, Thermo Fisher Scientific A-11055; anti-goat Alexa Fluor Plus 647, Thermo Fisher Scientific A32849; anti-Syrian hamster, Alexa Fluor 647, Thermo Fisher Scientific A18895.

Brain slices and optic and trigeminal nerves were then washed once in PBS with or without DAPI nuclear stain (1:50,000, Thermo Fisher) for 10 min, then washed twice for 10 min in PBS. After mounting, slices were imaged on a Zeiss LSM700 confocal microscope or Zeiss Airy980_MP (in confocal imaging mode).

### Light-sheet and confocal imaging of cleared mouse brain and cranial nerves

Immunocytochemistry performed on whole mouse brains (or sections thereof to allow them to fit into the light-sheet microscope tissue chamber) was as for brain tissue slices, except the incubation time in blocking buffer with both primary and secondary antibodies was 3 days at room temperature, and washes were performed with blocking buffer 3 times for an hour. Whole mouse brains or brains sectioned into several parts, and optic and trigeminal nerves, were then cleared for 2 weeks and 3 days, respectively at room temperature in CUBIC reagent R solution (distilled water containing 25% w/v N,N,N′,N′-tetrakis (2-hydroxypropyl)ethylenediamine), 15% w/v Triton X-100 and 25% w/v urea with a refractive index of 1.52). Brain sections imaged with a Zeiss LSZ1 light-sheet microscope and a 5x clearing lens were superglued to a custom-made sample holder and suspended from above in a tissue chamber containing CUBIC reagent R. Brain sections imaged with a Zeiss

Airy980_MP (in confocal imaging mode), with a Zeiss 25x lens set for imaging clearing solution with a refractive index of 1.52, were superglued to the base of a Petri dish filled with CUBIC reagent R solution.

### Analysis of tracer intensity

To measure the quantity of intracerebrally injected fluorescently labelled monomeric tau-441 exiting the brain over time via conduits within penetrating arteriole walls and around ascending venules, image stacks acquired during two-photon microscopy were analysed in ImageJ by measuring the average intensity of fluorescence for ROIs that either encompassed the entire arteriole or venule down to 50-100 μm, or in 10 or 20 μm substacks at different depths (Fig. 1j-n). Differences in the numbers of penetrating arterioles and ascending venules were corrected for (Fig.1o).

### Measurement of conduit diameter

Using Image J, conduit diameters were measured in confocal image stacks of brain tissue from 25 mice and 5 humans. A 5 pixel wide bar was placed perpendicularly across the conduit to measure diameter. When conduits appeared as “train tracks” (Fig. 4e), individual “rails” of the track were measured. Diameters of 10 conduits were measured (and averaged) per penetrating arteriole, ascending venule, pial artery and region of pial-arachnoid layers in mice, and 10 conduits per pial artery and region of pial-arachnoid layer in humans. In mice, a total of 10 penetrating arterioles (from 7 mice), 19 pial arteries (from 9 mice), 12 ascending venules (from 11 mice) and 22 regions of pial arachnoid-layers (from 15 mice) were measured. In humans, a total of 11 pial arteries (from 4 humans) and 12 regions of pial-arachnoid layers (from 5 humans) were measured. The averages per mouse or human were calculated and statistical tests applied using the mouse or human as the statistical unit.

### Software and Statistics

Image analysis was conducted using ImageJ software (NIH), Matlab (MathWorks), and IMARIS (v. 11.0, Oxford Instruments, Abingdon, UK). Data are presented as mean ± SEM, and obeyed normality (assessed with Shapiro-Wilk test). Comparisons of data were made using two-tailed Student *t* tests (corrected for multiple comparisons).

## Acknowledgements

Supported by: a Wellcome Clinical Fellowship to Ross Nortley (224632/Z/21/Z); a Sir Henry Wellcome fellowship to Harvey Davis (224049/Z/21/Z); an ERC Advanced Investigator Award (740427, BrainEnergy), a Wellcome Senior Investigator Award (219366/Z/19/Z) and an award from the BHF-UKDRI Centre for Vascular Dementia Research to David Attwell; and a CRUK Career Development Fellowship (CRUK-A19763), CRUK Senior Fellowship (RCCSCF-May22\100001) and CRUK CoL Centre support (CTRQQR-2021\100004) to Sophie Acton. We thank Veronika Lachina for illustration advice.

## Extended Data Figures

**Extended Data Fig. 1.**
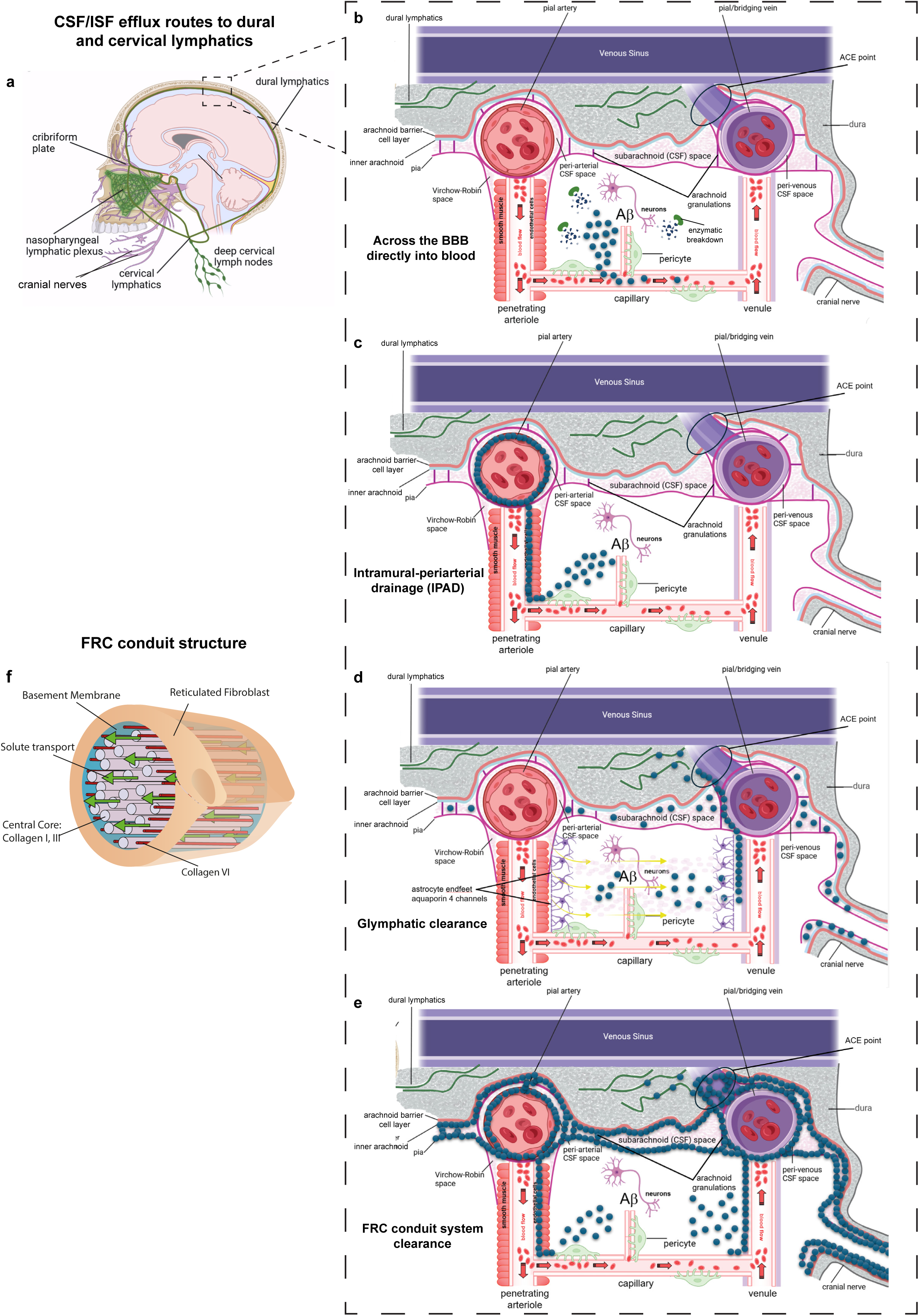
Schematic diagram of potential exit route from brain for proteins. **(a)** Routes for efflux of fluid and solutes from the brain via the dural lymphatics (green) which pass out of the brain posteriorly to the deep cervical lymph nodes, and may also pass out of the brain through the cribriform plate above the nose to reach the cervical lymph nodes. In the nasopharyngeal area there is a large lymphatic plexus. (**b**) Enlarged view of cortex, showing Aβ or tau produced within the parenchyma being removed by enzymatic degradation or by efflux across endothelial cells to the blood. (**c**) Parenchymal tau and Aβ can enter fibroblast reticular conduits at the arteriole end of the capillary bed, and ascend within conduits in the smooth muscle cell layer of penetrating arteriole walls, driven by arterial pulsation, to reach the pia (formerly known as the IPAD route for efflux). Here the proteins either continue in conduits around pial arterioles and arteries, and may exit the brain via conduits around arteries in cranial nerves, or they can traverse conduits linking arterioles and arteries to reach venules and veins, and from there leave the brain via bridging veins to reach the dural lymph. (**d**) Proteins can enter conduits around ascending venules, driven by water flow from the Virchow-Robin space around arterioles through astrocytes, and from there reach pial veins and exit the brain as above. This route was formerly attributed to a glymphatic system. (**e**) Efflux routes characterised in this paper, in FRC conduits within the wall of penetrating arterioles and around venules. (**f**) Schematic diagram of FRC wrapping a closed extracellular space that includes extracellular matrix proteins including collagen subtypes as well as extracellular space allowing fluid and solute movement.

**Extended Data Fig. 2.**
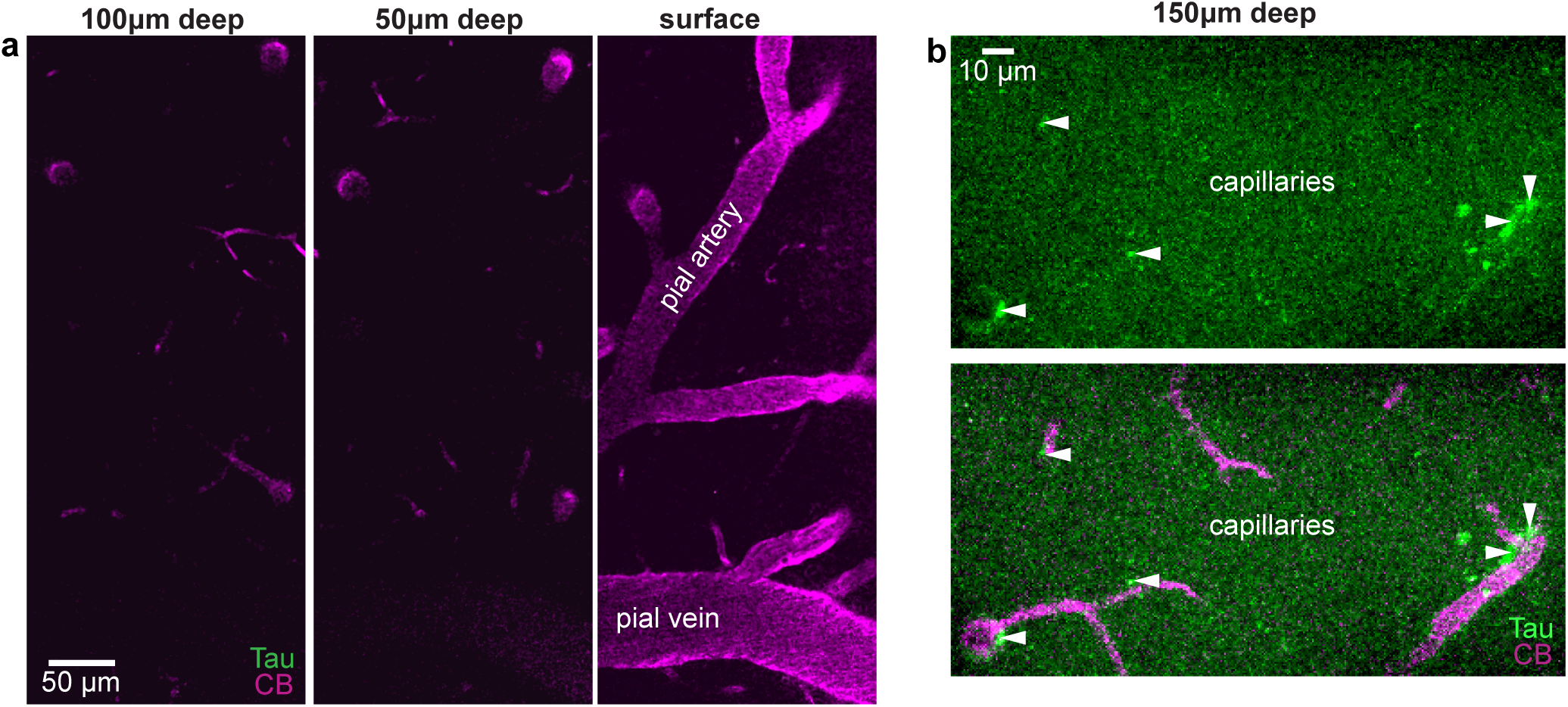
Tau moves through FRC conduits around capillaries in vivo. **(a)** Representative images at the brain surface and depths of 50 µm and 100 µm in same animal as Fig. 1a-c using in vivo two-photon-microscopy of 8-week-old WT mouse (via cranial window; CB = Cascade Blue (magenta) in blood) before intraparenchymal injection of fluorescently labelled (green) monomeric tau-441. **(b)** 5-min post intraparenchymal injection, fluorescent green monomeric tau-441 is seen diffusely within the brain parenchyma and concentrating around cortical capillaries (white arrowheads).

**Extended Data Fig 3.**
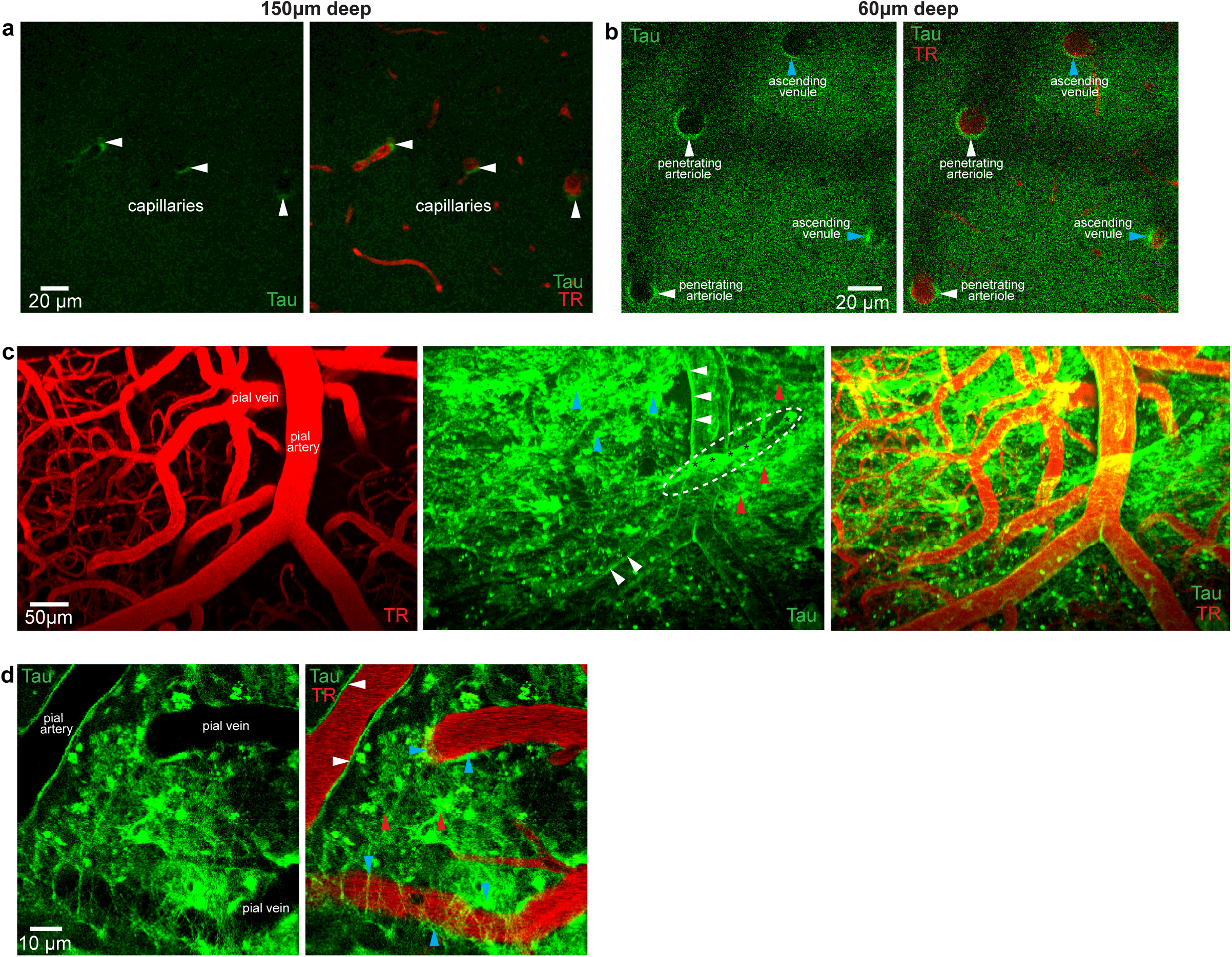
(a-d) Tau exits the brain in conduits around capillaries, penetrating arterioles and ascending venules, and pial vessels. Representative images of in vivo two-photon microscopy of 8-week-old WT mouse 30 min after topical application of fluorescently labelled (green) monomeric tau-441 via cranial window (TR = Texas Red). **(a)** Fluorescent tau-441 observed within the parenchyma and concentrating around the microvasculature at a depth of 150 µm. **(b)** Tau clearance from the cortical parenchyma within the walls of penetrating arterioles (white arrowheads) and around ascending venules (blue arrowheads) 60 µm below the brain surface. **(c)** Image stack of tau-441 clearing via FRC conduits within pial artery walls (white arrowheads), around pial veins (blue arrowheads) and the wider pial-arachnoid layers (red arrowheads) **(d)** At the pial surface, tau tracks through FRC conduits within pial arterial walls (white arrowheads), around pial veins (blue arrowheads) and in the wider pial-arachnoid mesh (red arrowheads).

**Extended Data Fig. 4.**
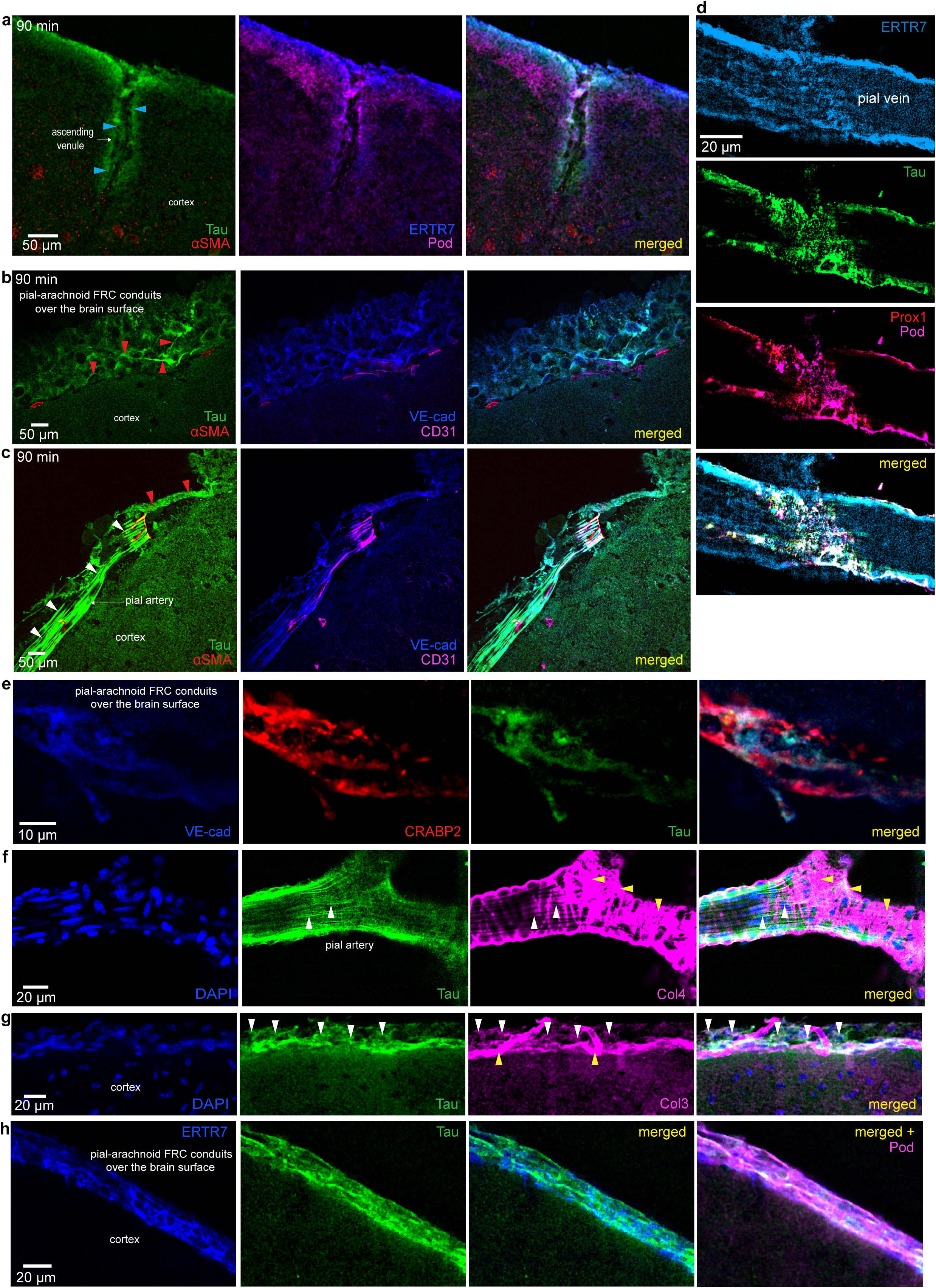
Exit of tau occurs via fibroblast reticular cell (FRC) conduits (following low volume ICM injection without cranial window formation). **(a-c)** Representative confocal images from 8-week-old WT mice 90 min following low-volume (500 nl) ICM injection of fluorescently labelled tau-441 (Tau, green), showing clearance via FRC conduits labelling for ERTR7 (collagen VI, blue) and podoplanin (Pod, magenta) surrounding an ascending venule (blue arrowheads, a), VE-cadherin labelled (VE-cad, blue) pial-arachnoid FRC conduits (red arrowheads, b) and within linear conduits in pial artery walls labelling for VE-cadherin (white arrowheads, c). **(d)** Tau clearance via pial-arachnoid FRC conduits around a pial vein, labelled for collagen VI (ERTR7, blue), Prox1 (red), and podoplanin (Pod, magenta). **(e-h)** Representative confocal images 90 min post ICM injection of fluorescently labelled (green) tau-441. **(e)** CRABP2 (red) and VE-Cadherin (blue) labelled fibroblasts of the inner-arachnoid layer of the pial-arachnoid conduits draining tau (green). **(f)** Tau draining via linear conduits within the pial artery wall which, along with the outer adventitia of the artery, label for collagen IV (Col4, magenta). **(g)** Tau clearance through pial-arachnoid FRC conduits labelling for collagen III (white arrowheads) (Col3, magenta, labels less strongly than does the pial vasculature (yellow arrowheads). **(h)** Collagen VI (ERTR7, blue) and Podoplanin (Pod, magenta) labelling highlight the mesh-like structure of the pial-arachnoid FRC conduits.

**Extended Data Fig. 5.**
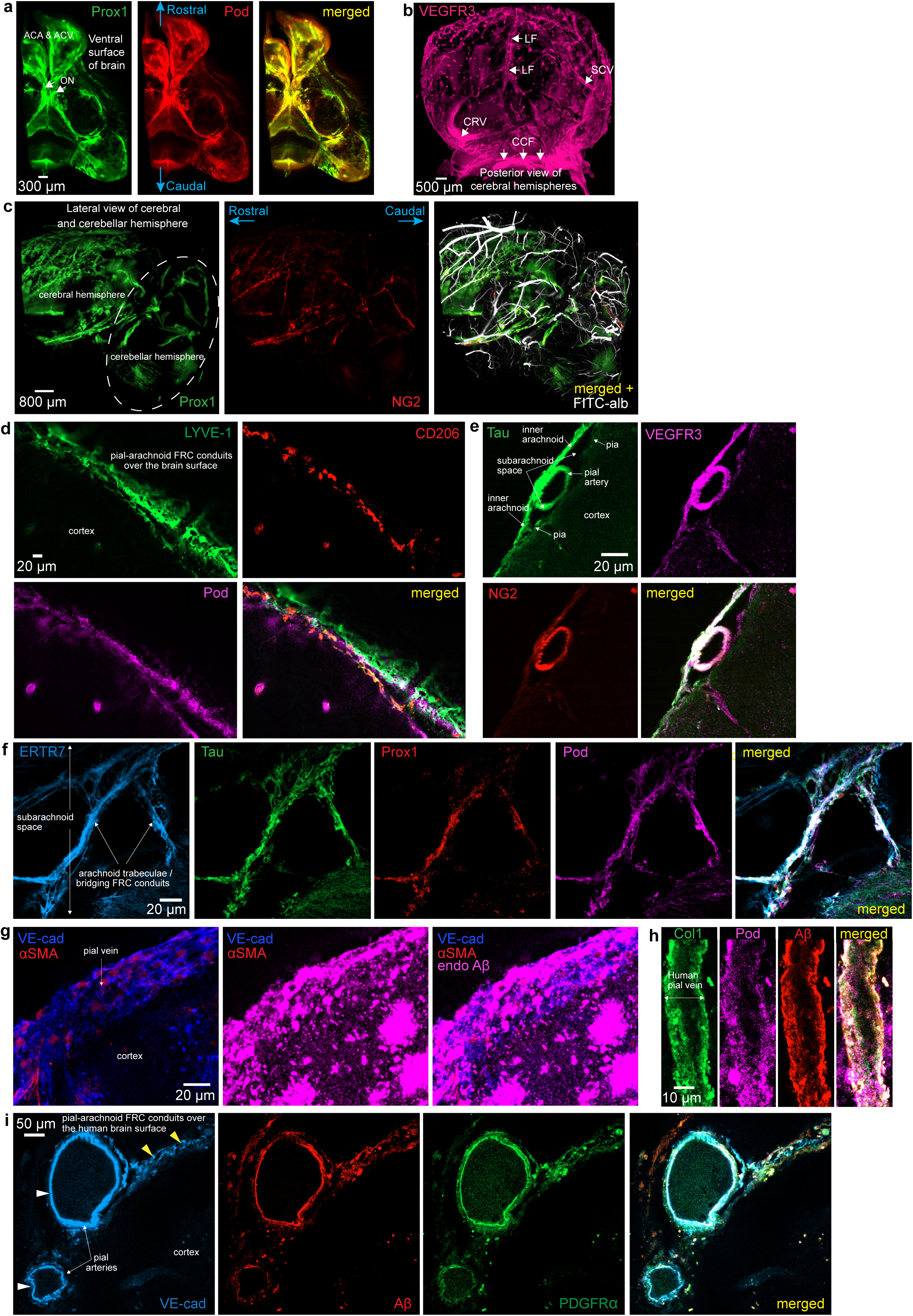
Further Imaging of FRC conduits over the ventral surface of the brain and of Aβ clearance in APP^NL-G-F^ mice and live human brain tissue. **(a)** Ventral surface of the cerebral hemispheres with colocalising Prox1 (green) and podoplanin immunolabelling of the pial-arachnoid FRC conduit network. Anterior cerebral artery (ACA), anterior cerebral vein region (ACV) and optic nerves (ON). **(b)** VEGFR3 labelling of pial-arachnoid FRC conduit mesh over the dorsal surface of the brain with prominence in regions around the longitudinal fissure (LF), and the surface of superior cerebral/pial veins (SCV). **(c)** Pial-arachnoid FRC conduits labelled for Prox1 (green) surface coverage of the posterior aspect of a cerebral and cerebellar hemisphere (dashed oval), with NG2 (red) labelling of pial arteries/arterioles and FITC-albumin (white) cast of the whole vasculature. **(d)** Representative confocal image from 8-week-old WT mouse demonstrating LYVE-1 (green) and podoplanin (pod, magenta) labelling of pial-arachnoid FRC conduits, and LYVE-1 (green) and CD206 labelling of perivascular microphages (red). **(e)** Clearance of tau via FRC conduits within a pial artery wall (SMA, red) in continuity with FRC conduits with the pial and inner arachnoid layers, labelled for VEGFR3 (magenta). (**f**) Tau can be observed tracking through FRC conduits labelling for collagen VI (ERTR7, blue), Prox1 (red) and podoplanin (Pod, magenta) within arachnoid trabeculae that span the CSF space of the basal cisterns. **(g)** Representative confocal image of endogenous Aβ (endoAβ, magenta) in an APP^NL-G-F^ mouse deposited as cortical plaques and throughout the pial-arachnoid FRC conduit mesh over the surface of a pial vein (with discontinuous smooth muscle cell labelling (α-SMA, Red). **(h-i)** Representative confocal images of fluorescently labelled (red) monomeric Aβ_1-42_ locating to conduits labelling for collagen I (Col1, green) and podoplanin (Pod, magenta) over the surface of a pial vein (yellow arrowheads, **h**) and VE-cadherin-labelled (VE-cad, blue) and PDGFRa (green) conduits (**i**) within the walls of pial arteries (white arrowheads) and pial-arachnoid conduits (yellow arrowheads) in live human tissue within 1 hour of incubation.

**Extended Data Table 1.**
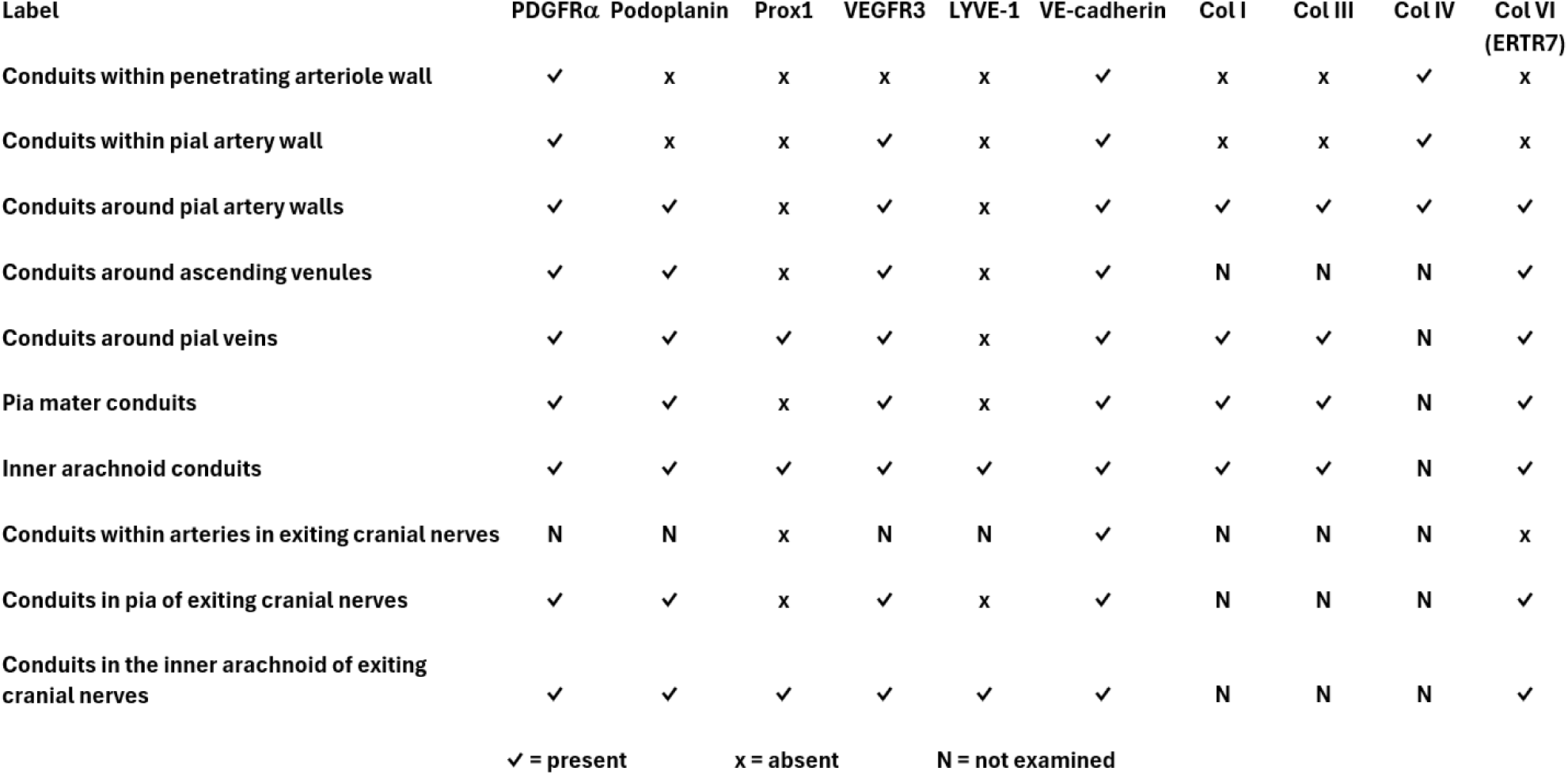
Markers labelling the FRC conduits at different locations.

